# Y2H-SCORES: A statistical framework to infer protein-protein interactions from next-generation yeast-two-hybrid sequence data

**DOI:** 10.1101/2020.09.08.288365

**Authors:** Valeria Velásquez-Zapata, J. Mitch Elmore, Sagnik Banerjee, Karin S. Dorman, Roger P. Wise

## Abstract

Interactomes embody one of the most effective representations of cellular behavior by revealing function through protein associations. In order to build these models at the organism scale, high-throughput techniques are required to identify interacting pairs of proteins. Next-generation interaction screening (NGIS) protocols that combine yeast two-hybrid (Y2H) with deep sequencing are promising approaches to generate protein-protein interaction networks in any organism. However, challenges remain to mining reliable information from these screens and thus, limit its broader implementation. Here, we describe a statistical framework, designated Y2H-SCORES, for analyzing high-throughput Y2H screens that considers key aspects of experimental design, normalization, and controls. Three quantitative ranking scores were implemented to identify interacting partners, comprising: **1)** significant enrichment under selection for positive interactions, **2)** degree of interaction specificity among multi-bait comparisons, and **3)** selection of *in-frame* interactors. Using simulation and an empirical dataset, we provide a quantitative assessment to predict interacting partners under a wide range of experimental scenarios, facilitating independent confirmation by one-to-one bait-prey tests. Simulation of Y2H-NGIS identified conditions that maximize detection of true interactors, which can be achieved with protocols such as prey library normalization, maintenance of larger culture volumes and replication of experimental treatments. Y2H-SCORES can be implemented in different yeast-based interaction screenings, accelerating the biological interpretation of experimental results. Proof-of-concept was demonstrated by discovery and validation of a novel interaction between the barley powdery mildew effector, AVR_A13_, with the vesicle-mediated thylakoid membrane biogenesis protein, HvTHF1.

**Author Summary:** Organisms respond to their environment through networks of interacting proteins and other biomolecules. In order to investigate these interacting proteins, many *in vitro* and *in vivo* techniques have been used. Among these, yeast two-hybrid (Y2H) has been integrated with next generation sequencing (NGS) to approach protein-protein interactions on a genome-wide scale. The fusion of these two methods has been termed next-generation-interaction screening, abbreviated as Y2H-NGIS. However, the massive and diverse data sets resulting from this technology have presented unique challenges to analysis. To address these challenges, we optimized the computational and statistical evaluation of Y2H-NGIS to provide metrics to identify high-confidence interacting proteins under a variety of dataset scenarios. Our proposed framework can be extended to different yeast-based interaction settings, utilizing the general principles of enrichment, specificity, and *in-frame* prey selection to accurately assemble protein-protein interaction networks. Lastly, we showed how the pipeline works experimentally, by identifying and validating a novel interaction between the barley powdery mildew effector AVR_A13_ and the barley vesicle-mediated thylakoid membrane biogenesis protein, HvTHF1. Y2H-SCORES software is available at GitHub repository https://github.com/Wiselab2/Y2H-SCORES.

## Introduction

Investigations into the molecular interactions among hosts and pathogens has benefited from the plethora of omics datasets that can be used for the prediction of gene and protein networks. For plants, this knowledge can be used to guide modern plant breeding efforts through the identification of resistance proteins and their co-functional partners (1, 2). In animal models, accurate networks can lead to the determination of drug targets and vaccines by providing a detailed view of the host-pathogen interactions that shape the host immune response (3–6).

The reconstruction of signaling networks is one of the most efficient methods to understand molecular events during host-pathogen interactions (7). These models can be depicted using protein interactome networks, where nodes represent proteins and edges represent physical interactions (8). The yeast two-hybrid (Y2H) screen is one of most powerful tools for uncovering new protein-protein interactions (PPI), discerning connections between bait and prey proteins while correcting for biases in their cell concentrations and affinity (9). However, traditional Y2H screens involve a labor-intensive step where distinct yeast colonies growing on selective media are picked, and Sanger-sequenced to identify prey cDNA fragments. Thus, to optimize genome-scale screens and obtain interactome data in a time-efficient manner, more recent approaches, collectively termed next-generation interaction screening (NGIS), use deep sequencing to score the result of Y2H screens. These innovations facilitate quantitative measures of bait-prey interactions, and importantly, do not require open-reading-frame sequence libraries from the organism(s) of interest (10–15).

Despite the methodological advantages of Y2H-NGIS, there remain overlooked informatics and statistical challenges. **1)** Most current pipelines map and quantify total reads while ignoring prey-fusion reads (reads containing both Y2H plasmid and prey cDNA) that provide frame information of the cDNA fusion protein. **2)** There is no consensus regarding what control(s) are more appropriate to signify the background interactivity of the preys and to help to distinguish true interactions (10–14, 16–18). **3)** Despite its importance, most existing methods do not assess data normalization, or implement inappropriate normalization methods for Y2H-NGIS data (11–14, 16, 18). High-throughput sequencing datasets, where read counts quantify signal strength, e.g., RNA-Seq, require normalization, as there are external factors, aside from experimental treatments, that influence read counts (19). Normalization methods, such as those used for RNA-Seq, assume that most genes in the sample are not differentially enriched (DE). However, in Y2H-NGIS experiments, the enrichment of each prey is determined by completely different factors under the two conditions. In non-selected conditions, the prey’s relative abundance in the library determines the enrichment, while under selection, it is the prey’s ability to activate the reporter, via interaction with the bait, or by auto-activation. Most if not all prey will therefore be DE in the selected condition. Finally, **4)** with no consensus on the appropriate data analysis, nor even how to report the results, whether ratios of counts, log fold-change from DE analysis, or a custom score function, it is nearly impossible to compare Y2H-NGIS studies, and there are reports that Y2H-NGIS-based interaction predictions suffer high false positive rates (10–14, 16–18). Hence, there is a need for robust and consistent statistical models that make use of all the available information in Y2H-NGIS data.

We optimized the protocol proposed by Pashkova and colleagues (11) to sub-culture diploid yeast populations that carry bait and prey plasmids under two batch conditions: **1)** diploid growth obtained in what we call the non-selected condition (SC-Leu-Trp), used as background level of library prey abundance and **2)** interaction test, or the selected condition (SC-Leu-Trp-His), which theoretically only allow the growth of diploid populations with positive bait-prey interactions through activation of the reporter gene. Diploids that grow under selection comprise a group of preys that can be split into true and false interactors. Here we define true interactors as those preys that can be verified as bait-specific in a binary Y2H interaction test with the appropriate negative controls. In contrast, false interactors represent the group of selected preys which do not produce a positive binary test, nor are bait-specific (*i.e*., preys that auto-activate the reporter). Previously, we developed a robust informatics pipeline, designated NGPINT, to identify candidate interacting partners obtained from Y2H-NGIS, including mapping reads to the reference prey genome, reconstruction of prey fragments, and distinguishing fusions with the prey activation domain (20). Here, we propose statistical methods to rank the resulting preys, distinguish false vs. true interactors, providing to the user a high-confidence list of candidates. This ranking system, designated Y2H-SCORES, is an aggregate of three experimental outcomes: **1)** the non-selected population as a baseline to detect which preys are significantly enriched under selection, **2)** selected samples as a control baseline to measure the specificity of a prey, and **3)** fusion read information to identify *in-frame* enrichment of the prey fragments under selection.

Using simulation and experimental validation we assessed the ability of Y2H-SCORES to successfully rank true vs. false interactions. Simulation of typical Y2H-NGIS data allowed us to demonstrate its robustness under different scenarios. Additionally, as a proof of concept, we used Y2H-SCORES to identify and confirm the interaction of the barley powdery mildew effector, AVR_A13_, with the barley vesicle-mediated thylakoid membrane biogenesis protein, HvTHF1. This interaction, accompanied with previous evidence (21) and expression quantitative trait loci (eQTL) associations (22), support the involvement of HvTHF1 with resistance to powdery mildew.

## Results

### The effect of normalization on Y2H-NGIS data

Principal component analysis (PCA) of the log-transformed Y2H-NGIS raw read counts identified selection as the major source of variability (Fig 1A). The effect of selection and baits (colors in Fig 1A) under selection are expected sources of variation, but we also expect all non-selected samples to resemble the cDNA used to build the prey library and the three replicates to cluster. Indeed, if there is more variation among replicates than baits, it could prove difficult to reproducibly identify bait interactors.

**Fig 1.**
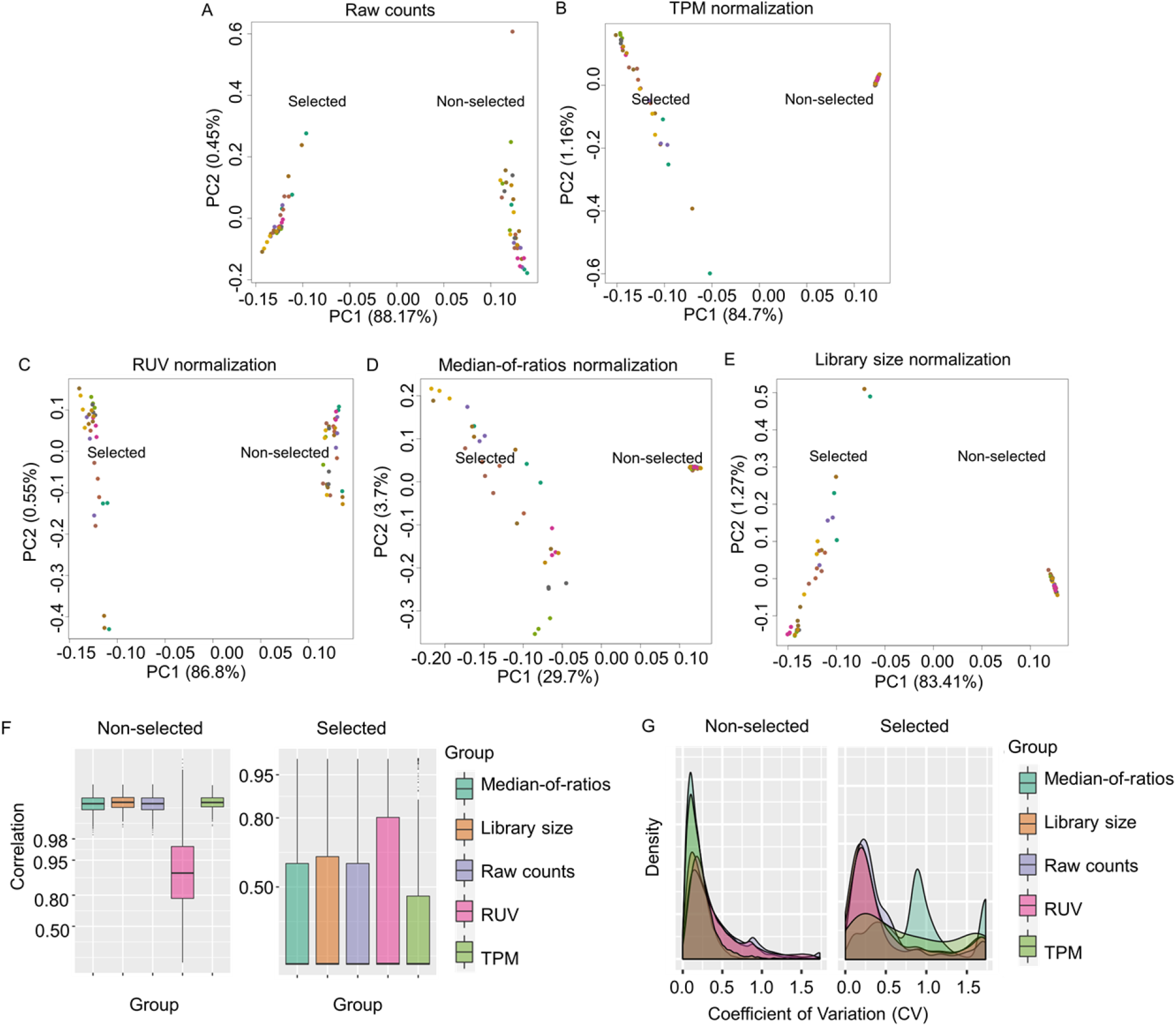
Effect of count normalization in Y2H-NGIS. A) PCA analysis of raw read counts and B) TPM, C) RUVs, D) Median-of-ratios and E) Library size normalized reads for selected (left) and non-selected (right) samples for 10 bait screenings (color coded). F) Boxplot of the pairwise correlation coefficients for raw and normalized read counts for all samples in non-selected and, separately, selected conditions. G) Coefficient of variation (CV) for each prey using different normalization methods in the bait AVR_A13_. Higher CV values may indicate poor performance because of a high variation between replicates.

We considered four normalization methods to reduce the experimental variation, particularly to reduce variation across replicates. All preys are expected to be differentially enriched (DE) in selected compared to non-selected samples; interactors should grow exponentially and non-interactors should not reproduce under selection. Our goal is to identify prey whose relative abundance in the selected samples increases over the relative abundance in the non-selected samples. Normalization methods appropriate for this goal include library size (23), transcripts (or in this case, prey fragments) per million (TPM) (24), and remove unwanted variation (RUVs) (25). Many normalization methods are designed to detect enrichment relative to unchanging reference genes, which simply do not exist in Y2H-NGIS data. Specifically, we used median-of-ratios (26), which assumes the majority of genes are not DE, as a control method that should fail to normalize Y2H-NGIS data. Fig 1B-E show the PCA plots of the total counts after implementation of the different normalization methods. TPM, RUVs, and library size reduced the variability in the non-selected samples to varying degrees but retained most of the other variation. The median-of-ratios method, in contrast, removed over half of the selected vs. non-selected variation. Thus, inappropriate normalization can eliminate important biological information that is used to infer interactors.

Ideally, all non-selected samples should resemble the prey library. Indeed, Pearson correlations of the count data between all pairs of non-selected samples exceeded 0.98 for all but RUVs normalization (Fig 1F), indicating that non-selected prey counts are largely bait-independent, as expected. In contrast, prey counts were much less correlated among selected samples, presumably reflecting the effect of the baits. Library size and TPM normalizations increased the non-selected sample correlation (Wilcoxon signed-rank test p-value of 1.88 x 10^-38^ and 1.12 x 10^-17^, respectively) over the raw counts (S1 Table), but RUVs normalization substantially decreased it (Wilcoxon signed-rank test p-value of 5.38 x 10^-81^). RUVs, which seeks factors explaining variation across replicates, may have retained greater variation within non-selected samples because we used just one factor to explain the technical variation.

Appropriate normalization should reduce inter-replicate variability. We measured the variability across replicates using the coefficient of variation (CV), computed for each prey. As illustrated in Fig 1G and S1 Fig, we found that RUVs, library size, and TPM normalizations reduced the CV compared to raw counts (Wilcoxon signed-rank test p-values <0.05, S2 Table). However, there is no one single method that works the best in all cases (S2 Table). For non-selected conditions, we found that all normalizations performed very well with CV peaks within 0 - 0.5, which indicate a low variation between replicates. TPM and median-of-ratios had the best performance with a narrower density. The median-of-ratios method can perform well in non-selected conditions since as compared with the selected conditions, they do not have any DE preys, resembling the transcriptome used to make the prey library (27). Contrast in performance was highest when normalizing selected samples. In this case, the ranking of methods was bait specific (S1 Fig, S2 Table), though the median-of-ratios consistently failed to reduce the variation. CV distributions for library size, TPM and RUVs peaked within 0 - 1, but the CV distribution for the median-of-ratios method peaked in the range 1 - 1.5 (Fig 1G). After evaluating the results from the Pearson correlation and CV analyses we decided to use library size normalization as the main method for the Y2H-NGIS dataset.

### Y2H-SCORES identifies true interacting partners

After optimizing normalization, we proposed a set of ranking scores based on statistical assessments of the count data to predict interacting partners. We considered the biological principles that define PPIs and key aspects of Y2H-NGIS, such as the experimental design, normalization, and controls (see methods). Summarizing, we modeled the total prey counts using a Negative Binomial (NB) regression and the *in-frame* fusion counts using the Binomial distribution. We created a modular set of three quantitative ranking scores, called Y2H-SCORES, to identify interacting partners: **1)** Enrichment score: a measure of significant enrichment under selection for positive interactions, using as control the non-selected samples; **2)** Specificity score: a measure of the specificity of a bait-prey interaction, using other selected baits as controls; and **3)** *In-frame* score: a measure of the enrichment for *in-frame* translational fusions in selected samples. To test Y2H-SCORES, we designed a Y2H-NGIS simulator, motivated by real data. The simulator includes true interactors (preys that are strongly selected only in the presence of their co-interacting bait), and auto-active/non-specific interactors. Auto-active preys activate the selection promoter without an interaction with the bait, while non-specific interactors survive selection because the product protein interacts with multiple baits (e.g., chaperones).

We began by simulating idealized conditions of 10 bait screens with three replicates, a cDNA prey library of 20,000 genes, 1 - 20 true interactors per bait, a stickiness factor (percentage of auto-active/non-specific preys in the library) of 0.1%, and a strength of true interactors above the 99.9^th^ percentile. The strength of true interactors was quantified with a fitness coefficient *e_ik_*, which we estimated for all prey in our real data. In this simulation, we reserved the top 0.1% of all estimated fitnesses for the true interactors, creating a sampling space that covers the maximum percentage of preys simulated from this group. This choice is based on experimental validations by library size: Pashova et al. (11) confirmed 8 out of ~15000 preys to be true interactors in their library, supported by our experiments which showed a similar trend, confirming between 1 and 25 in a ~36000 prey population.

We evaluated the performance of Y2H-SCORES using Receiver Operating Characteristic (ROC) and Precision Recall (PR) curves. ROC compares the true and false positive rates using different score value thresholds, while PR compares true and predicted positives (28). Fig 2A-C demonstrates that all scores performed well, separating true from auto-active/non-specific interactors. In this scenario all the scores achieved good performance: the enrichment, specificity and *in-frame* scores had a ROC Area Under the Curve (AUC) of 0.98 for enrichment, 1 for specificity and 0.99 for *in-frame*. The PR AUC was 0.47, 0.62, and 0.54, respectively. We plotted the PCA of the Y2H-SCORES under this scenario (Fig 2D) and we found the enrichment and specificity scores are more related with each other than with the *in-frame* score, nonetheless the three scores seem to provide different information based on their position in the plot.

**Fig 2.**
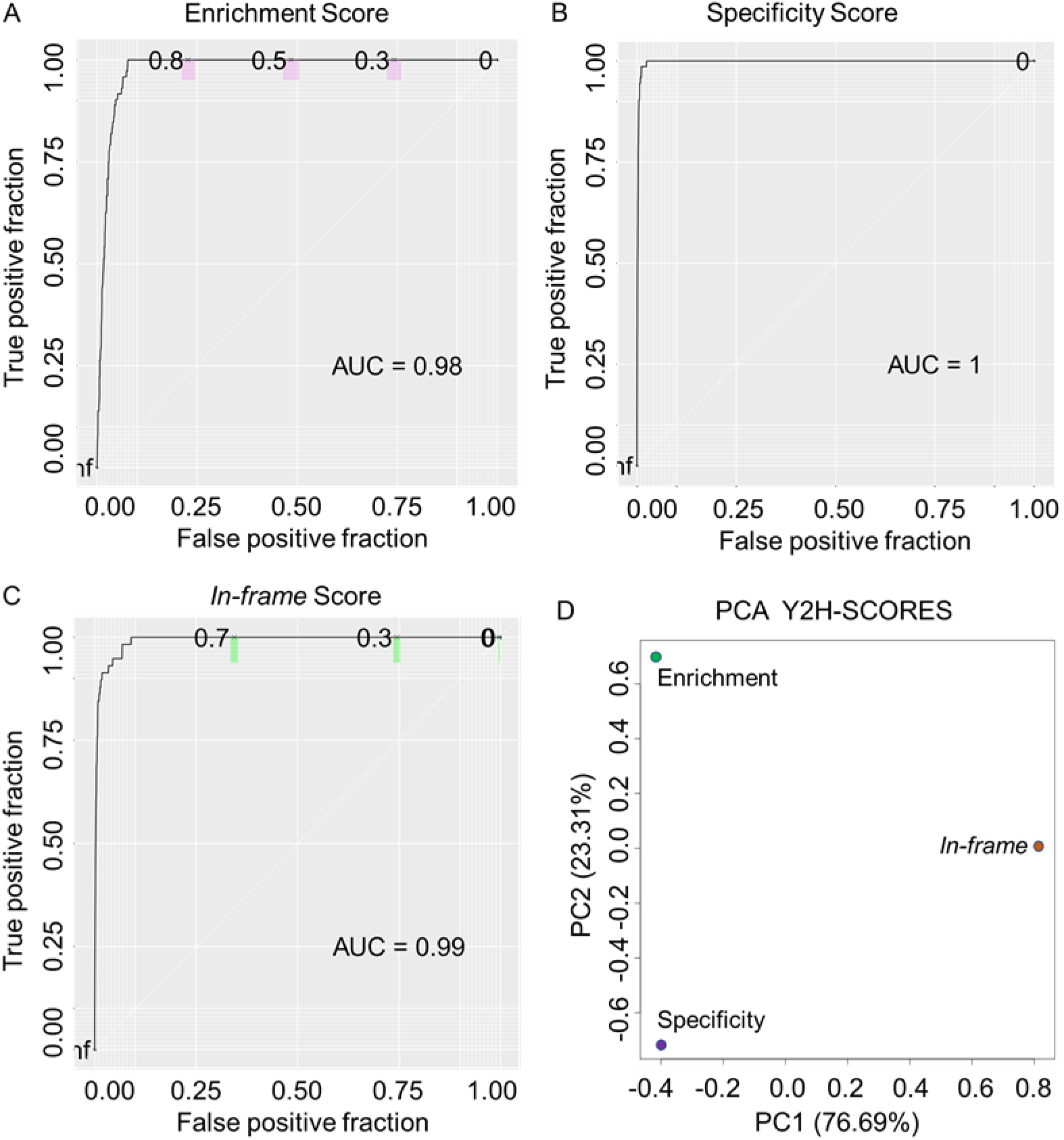
Performance of the ranking scores in an ideal scenario. A) to C) ROC curves of the enrichment, specificity, and *in-frame* scores. Colored sections represent 95% confidence intervals for the score values 0.7, 0.5, and 0.3. D) PCA of the Y2H-SCORES calculated under the ideal scenario.

### Y2H-SCORES overcomes challenging Y2H-NGIS scenarios

The ideal condition simulated for Fig 2 was discovered from an extensive simulation study where we explored the effect of several parameters that vary in experimental datasets, as defined in S3 Table. The simulator uses a Galton-Watson branching process followed by a NB model for generating total counts, and a binomial model for fusion counts (see S2 Text). To evaluate the performance of Y2H-SCORES, we varied the following parameters to simulate several Y2H-NGIS dataset scenarios: **1)** size of the prey library, to assess scalability; **2)** stickiness (*i.e*., the percentage of auto-active/non-specific preys in the library) and **3)** strength of true interactors, to vary the signal-to-noise ratio; **4)** overdispersion, to assess increasing levels of biological and experimental variation; **5)** proportion of true interactors in the prey library; to assess the role of genetic drift; **6)** number of baits and **7)** replicates, to assess power. Aside from the true interaction strength and the stickiness, these parameters are directly associated with the cDNA prey library and experimental conditions, which can be improved by the researcher. A graphical summary of the results is shown in Fig 3. Briefly, the scores were able to correctly identify true interactors even in extreme conditions, but the variation in performance helps us identify ideal experimental setting for detecting these interactors.

**Fig 3.**
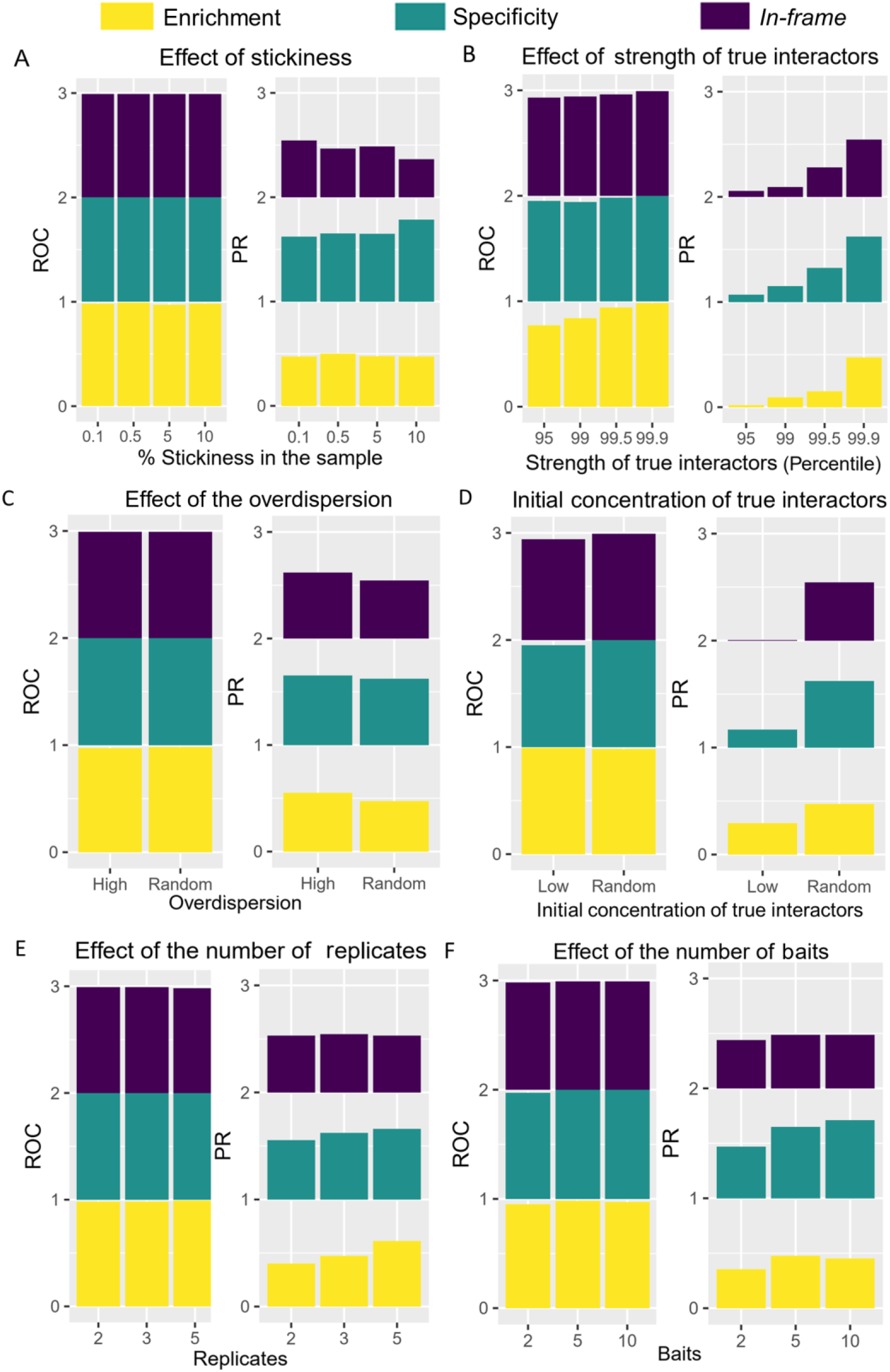
Effect of changes in the parameters that define Y2H-NGIS simulation. Examples of challenging scenarios were simulated to determine the Y2H-SCORES classification power. A) Stickiness (percentage of auto-active/non-specific preys in the library), B) Strength of true interactors, C) Overdispersion, D) Concentration of true interactors in the prey library, E) Number of replicates, and F) Number of baits. ROC and PR AUC values were reported for the enrichment, specificity, and *in-frame* scores.

The scalability of the Y2H-SCORES was evaluated by testing three prey library sizes (8000, 20000 and 40000 preys). We found that increasing the library size maintained the performance of the scores (S3 Table). The PR AUC values of the enrichment score oscillated between 0.47 and 0.68, the specificity from 0.62 and 0.80 and the *in-frame* score from 0.54 to 0.76, while the ROC AUC remained constant. This result shows that even with large library sizes Y2H-SCORES performs well and therefore, can still be used to identify protein-protein interactions.

We then tested the effect of the stickiness of the samples and the strength of true interactors on the Y2H-SCORES performance. The results from our simulations, shown in Fig 3A-B, suggest that Y2H-SCORES performance is less influenced by changes in the stickiness than by the strength of true interactors. Keeping the strength of true interactors above the 99.9 percentile and variations of the stickiness between 0.1% and 10%, did not cause major changes in the ROC and PR AUC values. This result indicates that Y2H-SCORES is able to identify auto-active/non-specific interactors, even in when they comprise 10% of the preys in the sample. In contrast, the strength of true interactors had a greater effect on the performance of Y2H-SCORES. As we decreased the strength of true interactors from the 99.9^th^ to the 95^th^ percentile, we found that the PR AUC values dropped to near zero. ROC curves were more stable, showing a gradual decrease. As expected, decreasing the signal-to-noise ratio in the system reduced the performance of Y2H-SCORES.

To evaluate the effect of experimental variation we tested changes in the overdispersion. We simulated two scenarios, either a high or random overdispersion in both the selected and non-selected condition. After estimating the overdispersion parameters observed in real data, we jointly sampled the proportion of preys and the overdispersion *φ_kN_* in the non-selected samples, and the fitness and overdispersion values *φ_iks_* in the selected samples, from the joint empirical distributions. In the overdispersed scenario, we resampled *φ_kN_* and *φ_iks_* values higher than the 90^th^ percentile of their densities (2.27 < *φ_kN_* < 13.42, 0.33 < *φ_iks_* < 2). The scores’ performance was maintained in scenarios with high overdispersion as measured by both PR and ROC AUC values (Fig 3C).

The initial proportion of each prey before culture expansion depends on the composition of the cDNA prey library, which can be controlled through experimental library normalization (29). We assumed the post-expansion prey proportions in the non-selected samples were identical to the unobserved prey proportions at the beginning of selection. Thus, we sampled these initial proportions from the observed non-selected proportions. We expect more inter-replicate variability and lower power to detect true interactors when the initial true interactor proportion is low because of initial sampling variation and greater genetic drift during culture growth. To simulate the effect of a low concentration of preys in the library we used the minimum proportion *q_ik_* that we observed in our experimental dataset, ~1 x 10^-8^, as reference value and assigned it to the true interactors in the “Low” condition (Fig 3D). Results from this analysis showed that the PR AUC decreased for all three scores. The enrichment and specificity scores decreased from 0.47 to 0.30, and from 0.62 to 0.17, respectively. The largest decrease was observed for the *in-frame* score, going from 0.54 to 0.005. Scenarios with low proportions of true interactors in the prey library caused a low number of total prey reads for that group in the non-selected condition, and a reduction or even absence of fusion reads (which normally represent a small fraction of the total number of reads). This trend was also observed in experimental datasets, where we observed a large number of preys with no fusion reads available in the non-selected samples.

Detection of DE prey with statistical confidence requires replication, but more replicates increase the time and cost of the sequencing project. We evaluated the effect of having two, three and five replicates. Increasing the number of replicates increased the performance of the enrichment and the specificity scores, while the *in-frame* score was not affected (Fig 3E, S3 Table). The *in-frame* score maintained a good performance even in cases with two replicates, with PR AUC around 0.53, but the enrichment and specificity scores had reduced performance. The enrichment score had the greatest reduction in the PR AUC values going from 0.61 (five replicates) to 0.40 (two replicates), and the specificity PR AUC values went from 0.65 to 0.55. Finally, we tested the effect of the number of baits using values from two to ten (Fig 3F). The enrichment and *in-frame* scores showed a decrease in their PR AUC values only in the case with two replicates with values of 0.35 and 0.44, respectively. In contrast, the performance of the specificity score improved with more baits in the simulation, with PR AUC values increasing from 0.47 to 0.71. The specificity information provided by the additional bait screenings increases the resolving power of the specificity score.

### Y2H-SCORES discards auto-active preys and identifies a new interaction between AVR_A13_ and HvTHF1

As proof of concept, we used Y2H-SCORES to identify interactors of the AVR_A13_ effector from the barley powdery mildew pathogen, *Blumeria graminis* f. sp. *hordei (Bgh)* (30). Our framework allowed us to discard auto-active interactors and to confirm the interaction between AVR_A13_ and the barley vesicle-mediated thylakoid membrane biogenesis protein, HvTHF1. We screened AVR_A13_ using an experimental setting of three replicates in the non-selected and selected condition. We ran our bioinformatic pipeline to map and quantify reads in the samples and obtained the prey regions for cloning. After this, we calculated the three Y2H-SCORES and created a Borda ensemble (31) to obtain a list of candidate interactors. Interestingly, after running Y2H-SCORES with different normalizations we found an increase in the number of highly ranked candidates with the median-of-ratios method. When we dissected this trend, we observed an increase in the number of candidates with high enrichment and *in-frame* scores and low specificity score (Wilcoxon ranked-sum test, S4 Table; S2 Fig), theoretically indicating auto-active/non-specific preys. The top-scoring preys unique to this list had low specificity scores across all normalization methods (S5 Table). We performed binary Y2H with two of these preys, corresponding to the gene IDs HORVU2Hr1G060120 (TCP family transcription factor 4) and HORVU2Hr1G024160 (Chaperone protein DnaJ-related), and confirmed that they were auto-active (S3 Fig), as they interacted with empty vector and luciferase (non-native protein).

Using Y2H-SCORES calculated from library size normalization, we focused on preys with high Borda ensemble scores, and therefore, high Y2H-SCORES values. HvTHF1, corresponding to the barley gene ID HORVU2Hr1G041260, was identified as the top candidate interactor of AVR_A13_ with the following Y2H-SCORES: enrichment of 0.9743, specificity of 0.8986, *in-frame* of 1 and Borda ensemble of 1203.73. Fig 4A shows the IGV alignment of this prey, depicting part of its fusion reads. To test the binary interaction, the *AVR_a13_* bait sequence was fused with the GAL4 transcription factor binding domain (GAL4-BD), while the *HvThf1* prey sequence was fused with the GAL4 DNA activation domain (GAL4-AD). After mating, diploid selection in SC-LW media and selection of the interaction with SC-LWH and SC-LWH + 3AT, we obtained a strong and specific positive interaction. Fig 4B shows the results from this binary test, validating the interaction between this protein and the effector.

**Fig 4.**
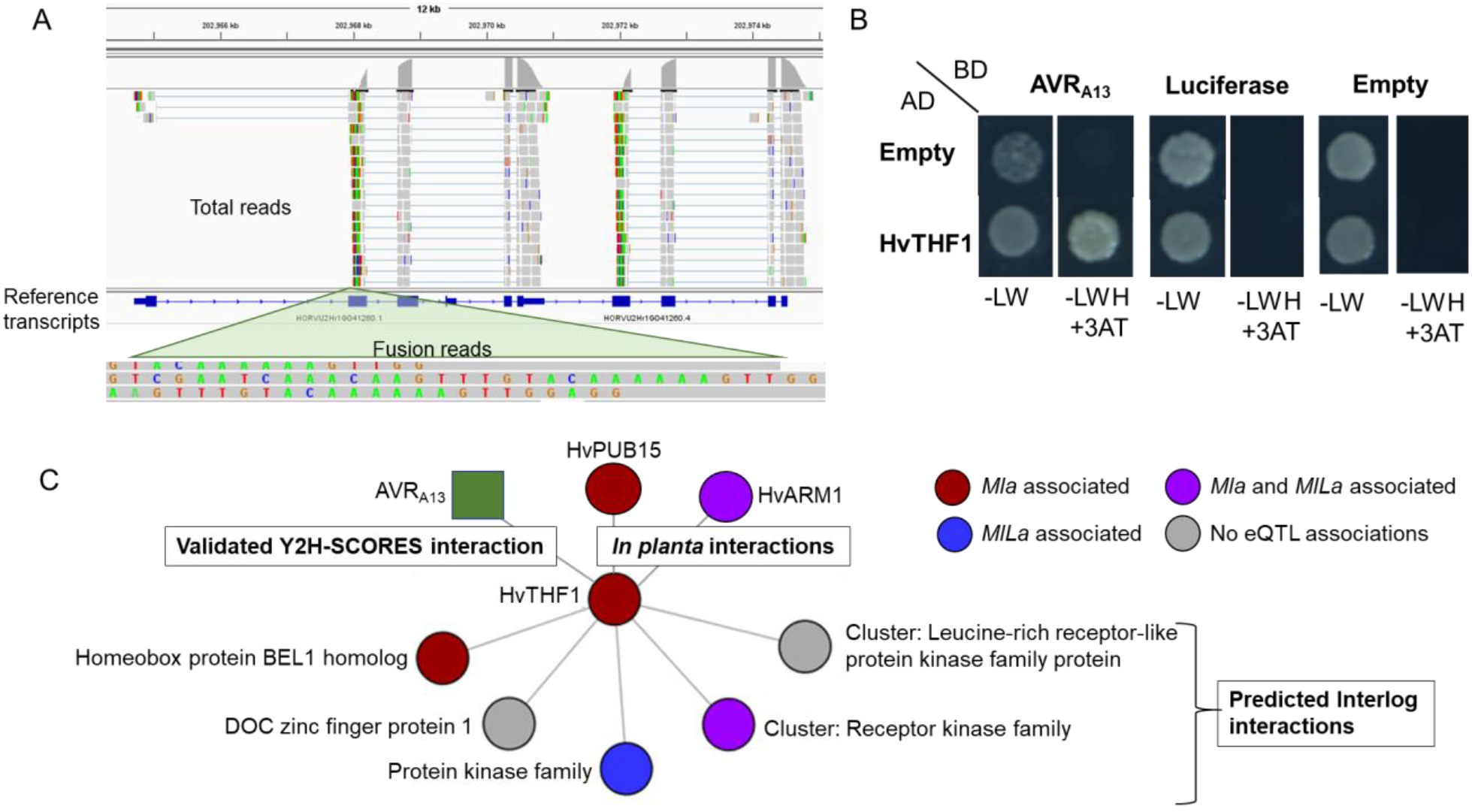
Experimental validation of the interaction between the barley powdery mildew effector, AVR_A13_ and the vesicle-mediated thylakoid membrane biogenesis protein, HvTHF1. A) IGV mapping of reads to the prey region and zoom to fusion reads, used for primer design and cloning. B) Binary Y2H test between AVR_A13_ and HvTHF1 showing the diploids control (SC-LW), the stringent interaction (SC-LWH+3AT), and tests with luciferase and empty-bait vector to show the specificity of the interaction. C) Prediction of interlogs for the AVR_A13_ effector target, HvTHF1 (validated Y2H-SCORES interaction). HvPUB15 and HvARM1 are shown as *in planta* interactors of HvTHF1 (21). *Trans*-eQTL associations (22) with the *Mla1* (*mildew resistance locus a1*) and *MlLa (Laevigatum* resistance locus) are color coded.

The prey consisted of a cDNA fragment of 817 nucleotides with an open reading frame of 162 amino acids (S1 Text contains fasta files with the prey sequence and protein translation). We analyzed the prey fragment peptide to identify the interacting domains and we found it contains a thylakoid formation protein domain. To position HvTHF1 in a signaling context we predicted protein-protein interactors using interlogs (32–35). As illustrated in Fig 4C, we found 18 predicted interactions with four main protein families: Homeobox protein BEL1, DOF zinc finger protein, protein kinase, receptor kinase and leucin-rich receptor-like kinase. Interestingly, previous expression quantitative trait locus (eQTL) analysis indicated that *Thf1* associates significantly with the *Mla1* (*mildew resistance locus a1*) *trans*-eQTL (22). Many of the predicted interactors for the THF1 protein are also associated with *trans*-eQTL at different powdery mildew infection stages, *MlLa (Laevigatum* resistance locus) at penetration and *Mla1* during haustorial development (22).

## Discussion

### Analysis of Y2H-NGIS data

Analysis of Y2H-NGIS data is challenging due to the complexity of the raw dataset (composed of total and fusion prey reads under both selective and non-selective conditions) and the substantial variability across replicates. This variability may be due to different factors in the experiments, including stochastic mating, genetic drift, cell viability and composition of the prey library aliquots. Banerjee and colleagues (20) proposed a bioinformatic pipeline to map and quantify raw reads from Y2H-NGIS, providing both total and fusion prey counts under selective and non-selective conditions. In this report, we outline Y2H-SCORES, a framework to rank candidate prey/bait interactions based on these count data. Using a Negative Binomial (NB) regression we modeled total counts, and the *in-frame* fusion counts were analyzed using the Binomial distribution. From these models we designed Y2H-SCORES to identify interacting partners based on their properties: **1)** Enrichment score: measures the enrichment under selection for positive interactions, as compared with non-selected conditions; **2)** Specificity score: assigns higher values to unique bait-prey interactions, as compared to prey selected in multiple bait screens; and **3)** *In-frame* score: measures the enrichment for *in-frame* proteins in selected samples. We validated the method and used simulation to evaluate the impact of several experimental factors on the power to detect true interactions and the accuracy of the rankings. We found that normalization methods and controls have a profound impact on the amount of information that can be used to identify interactors.

Normalization significantly modified the variation among replicates. Utilizing methods whose assumptions are satisfied by the Y2H-NGIS dataset leads to a more successful interpretation of the observed variation to infer protein interactors. Library size, TPM and RUVs are appropriate normalization methods for Y2H-NGIS data, but their ability to reduce variance within replicates varied, therefore, we recommend that users evaluate them individually and decide which one works better for their experiment. Median-of-ratios normalization, commonly used for RNA-seq data, is not appropriate for Y2H-NGIS data, and applying this method increased the number of candidate interactors with high enrichment and *in-frame* scores and low specificity score, compared to the other three normalization methods. Median-of-ratios normalization appears to promote non-interactors and auto-active/non-specific preys (which activate the selection promoter without an interaction with the bait, or interact with multiple baits), since the top-scoring preys unique to this list had low specificity scores across all normalization methods (S5 Table) and two of them were confirmed as auto-active (S3 Fig). Performing appropriate normalizations removed these auto-active preys from the top ranked list. Overall, median-or-ratios normalization produced lists with higher enrichment and *in-frame* scores, but lower specificity scores than the other methods (Wilcoxon rank-sum test, S4 Table).

Additionally, different controls that have been proposed for Y2H-NGIS can be used to measure different properties of an interactor. First, we used non-selected controls as an enrichment baseline of preys, allowing the implementation of the enrichment and *in-frame* scores. Second, we demonstrated the advantage of using screenings under selection for multiple baits as a second type of control that provides information for the specificity score. The baits used for this purpose may contain an empty bait, a non-native bait, or a set of baits of interest. If a combination of these baits is used in the experiment, auto-active preys should have lower specificity scores relative to non-specific preys, providing some separation. Our PCA analysis of Y2H-SCORES (Fig 2D) suggest independent information coming from each of the three scores, hence we recommend using non-selected and multiple selected bait controls to allow the implementation of all three scores to obtain a high-confidence list of interactors. The modularity of Y2H-SCORES also allows for partial score calculations, which will depend on the type of control used.

Development and testing of Y2H-SCORES has yielded some suggestions for the design of Y2H-NGIS experiments. Calculating the enrichment and the *in-frame* scores requires selected and non-selected samples. As we demonstrated (Fig 1F), composition of the non-selected sample is almost identical regardless of the bait (Pearson correlation > 0.98), information that can be utilized to reduce the number of sequenced samples, e.g., using a few random baits as non-selected controls for multiple baits in the same mating design. In contrast, the specificity score requires at least two different bait screenings, with better results as the number and type is increased. This strategy exploits the count information of multiple selected baits to identify auto-active/non-specific preys, giving priority to specific candidate interactors. However, researchers are often interested in a particular biological process and might screen several baits involved in a specific signaling pathway. In this case, preys that interact with multiple baits and exhibit low specificity might be prioritized for downstream validation. In that case, we also recommend using empty and/or non-native controls to discard auto-active preys. Thus, depending on experimental goals, the specificity score can be leveraged to find novel co-interacting partners of multiple proteins of interest.

### Experimental setting and optimization of Y2H-SCORES

Different experiment scenarios allowed us to test the robustness of Y2H-SCORES and identify the most challenging dataset types (Fig 3). The three scores were affected differently depending on the simulation scenario. Scenarios with low strength and low concentration of true interactors in the prey library imposed the most challenging conditions for these scores, reducing the ROC and PR AUC values. The enrichment score was more affected by the strength of true interactors while the *in-frame* score was more affected by the concentration of true interactors in the prey library, due to the inherent lower number of fusion reads. This analysis led us to explore aspects of experimental design that could be adjusted to increase the accuracy and sensitivity of interactor detection via Y2H-NGIS. These include the experimental prey library normalization, the number of replicates and baits in the experiment, sequencing depth, and scaling of the experiment setting.

First, library normalization can optimize the proportion of each prey in the library, in the non-selected samples and at the start of selection. In a typical cDNA library, the relative abundance of species derived from different genes can span many orders of magnitude. Normalizing the prey library to reduce high-abundance cDNAs reduces the stochasticity and noise in prey counts. After normalizing the prey library, experimentalists should also ensure the number of yeast recipient cells are sufficient to represent such library in the screens, for which they can use procedures as described by Krishnamani et al. (36). Our simulations found that low abundance interactor preys in the library (200 times lower than the expected prey abundance) can be detected in Y2H-NGIS. However, as the initial concentration of a true interactor decreases, a stronger affinity for the bait (relative to the interaction strength of auto-active/non-specific preys) is required for reliable detection. Thus, normalizing the prey library can reveal weaker true interactors.

A second parameter, the number of replicates in Y2H-NGIS, represents a cost-power trade off as in most “omics” experiments. For small numbers of replicates, as tested in the simulations, we found at least three replicates of Y2H-NGIS were needed to maintain the performance of the three scores. As expected, we observed better results as the number of replicates increased, especially for the enrichment score. We did not test replicate numbers greater than 5 since this does not represent typical experimental practices, and if one had to choose, increasing the number of baits would yield more biological information, since it would increase the performance of the specificity score. We anticipate that increasing the number of replicates would decrease the false discovery rate as it is reported for techniques such as RNA-Seq (37). Controlling for false discovery rate also informed our selection of DESeq2 as the tool for calculating differential enrichment due to the documented outperformance in low replicate numbers (19, 37).

It has been reported that reducing overdispersion in counts can improve sensitivity and accuracy of Y2H-NGIS (11). We did not observe a decrease in the performance of the Y2H-SCORES as we increased the overdispersion, which may be explained by the strategies that we took in our experiments to control it. As a result, the overdispersions estimated from our data may already be lower than what we could have observed with a different experimental setting. The main recommended strategies to control overdispersion among samples include maintaining a large-scale mating (in our experiment, 1.8 x 10^8^ bait and 5 x 10^7^ prey cells, resulting in ~2 x 10^9^ diploid cells) and subsequent high culture volumes of non-selected and selected samples (typically 800 ml per sample in 2-liter fluted Erlenmeyer flasks for ca. 36,000 preys in the library). Increasing the volume reduces the stochasticity of the prey population before mating and during culture expansion. Stochasticity is most notable in selected growth, where genetic drift dominates as the viable prey population shrinks. Population bottlenecks must be avoided throughout the experiment, which implies increasing the aliquot size in every culture step, including the final sampling for sequencing, as reported by Pashkova and associates (11). We also recommend adjusting the sequencing depth to match the prey library size and specifically increasing depth for the more complex, non-selected samples. Having a high depth in non-selected samples also increases the number of fusion reads, the major challenge for the successful implementation of the *in-frame* score in our simulations.

### Extrapolation of Y2H-SCORES to other yeast-based interaction methods

The analytical framework we propose here can be extrapolated to other yeast-based interaction settings, e.g., testing for DNA-protein interactions through yeast one-hybrid (38) or multiplexed yeast two-hybrid (12). The similarities and limitations among these techniques make them suitable candidates for the implementation of the Y2H-SCORES framework. All share the same true interactor properties: they should be enriched in selected samples, be specific to a bait screen and be selected *in-frame*. In addition, these techniques require control for false positives with appropriate statistics. Currently, these techniques propose a solid media selection of interactors, nonetheless the batch culture experimental setting proposed by Pashkova and colleagues (11) can also be applied to these contexts, increasing the reproducibility and facilitating the experimental workflow. Once output counts are obtained, it will be possible to calculate the Y2H-SCORES and use them to identify true interactors.

### Experimental validation of Y2H-SCORES identifies a new interaction between AVR_A13_ and HvTHF1

As an example of the effectiveness of our ranking scores, we identified and validated a new interaction between the barley powdery mildew effector, AVR_A13_, and the vesicle-mediated thylakoid membrane biogenesis protein, HvTHF1. HvTHF1 has been found to interact in yeast and *in planta* with proteins encoded by the U-box/armadillo-repeat E3 ligase *HvPUB15* and a partial gene duplicate, *HvARM1* (for *H. vulgare* Armadillo 1) (21). Neo-functionalization of *HvARM1* increases resistance to powdery mildew and provides a link between plastid function and colonization by biotrophic pathogens. In the broader context of plant-pathogen interactions, THF1 and its homologs interact with different protein mediators of plant resistance and susceptibility. The wheat homolog of *HvThf1, TaToxABP1*, encodes a target of the necrotizing Toxin A effector from the tan-spot fungal pathogen, *Pyrenophora tritici-repentis*, which has been associated with ROS (reactive oxygen species) burst (39, 40). In addition, THF1 also has been found to destabilize several NB-LRR-type resistance proteins by binding their I2-like coiled-coil (CC) domains (41). Previous eQTL analysis performed by our group, combined with an interlogs search, revealed significant associations of *Thf1* and its primary interactors with the *Mla1* and *MlLa trans*-eQTL (22). These *trans*-eQTL associations (Fig 4C) were observed at different powdery mildew infection stages, *MlLa* at penetration, and *Mla1* during haustorial development. These results provide an additional entry point to support the involvement of HvTHF1 with resistance to powdery mildew disease.

## Methods

### Normalization

We implemented library size (23), transcripts per million (TPM) (24), removing unwanted variation (RUVs) with replicate control samples (25) and the median-of-ratios (26) normalization methods. RUVs was applied to selected and non-selected samples separately, using k=1 factor and grouping the three replicates for each bait in the selected condition, and all replicates for all baits in the non-selected condition. Median-of-ratios was applied for all baits and conditions, grouping the replicates for each bait-condition combination. The coefficient of variation (CV) for each prey was calculated analyzing each bait-condition combination separately and grouping the three replicates for each prey. Pairwise differences between the distributions of the CV were detected with a Wilcoxon signed-rank test on the prey CVs computed after application of each normalization method for each bait. Pearson correlation was calculated for each method separately and within replicates for each bait-condition combination. Correlation differences between each normalization method were assessed using a Wilcoxon signed-rank test.

### Differential enrichment analysis of prey counts

We modeled the optionally normalized prey count data with Negative Binomial (NB) regression (42). This distribution allows for the effects of selection on the mean counts, while accounting for overdispersion across replicates, a consequence of biological and experimental variability. We calculated the significance and magnitude of the enrichment for each prey gene interacting with each bait using the DESeq2 model (42). Two different baseline controls were used to identify interactors: the non-selected condition for enrichment and the other selected baits for specificity.

### Modeling fusion reads

Let *Y_ikc_* be the number of *in-frame* reads out of a total *F_ikc_* fusion reads for prey *k* mating with bait *i* in condition *c* = *N,S* (*N*=non-selected, *S*=selected). We modeled *Y_ikc_ ~ Bin*(*F_ikc_, π_ikc_*), where *π_ikc_* is the proportion of *in-frame* reads. To test *in-frame* enrichment under selection, we pose the hypothesis: *H_o_*:*π_ikN_* = *π_iks_* vs. *H_a_*:*π_ikN_* < *π_iks_*, testing for an increase in the *in-frame* read proportion under selection. We evaluated this hypothesis using the *Z*-score statistic *ρ_ik_*:

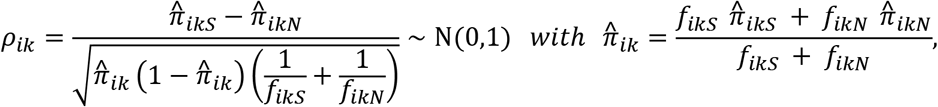

where 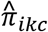 is the observed *in-frame* read proportion and *f_ikc_* is the observed number of fusion reads for prey *k* mated with bait *i* in condition *c*.

### Y2H-SCORES

We implemented and validated a ranking score system, designated Y2H-SCORES, for identifying interacting partners from Y2H-NGIS. It is comprised of three elements, each in the range [0,1], with values close to 1 indicating high support for a true interaction. Throughout, *np* is the number of prey and *nb* is the number of baits.

#### Enrichment score

This score quantifies the level of enrichment of a prey *k* under selection with bait *i* relative to non-selection. Let *p_ik_* be the *p*-value and *f_ik_* the log2 fold-change in the normalized counts of prey *k* interacting with bait *i* in selected over non-selected condition as given by DESeq2. The score consists of a system of ranks of log2 fold-change within ranks of *p*-values to prioritize interactors. We first define the rank for *p_ik_*: Consider *G_α_* = {(*i, k*): 1 ≤ *i* ≤ *n_b_*, 1 ≤ *k* ≤ *n_p_, p_ik_* ≤ *α*}, the set of putatively interacting prey/bait combinations with *p*-values *p_ik_* ≤ *α*. Bait/prey combinations with *p*-values larger than *α* are assigned an enrichment score of zero. The *p*-value enrichment score for the remaining bait/prey combinations is:

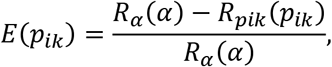

where *R_α_*(*p*) is the rank of *p*-value *p* among the *p_ik_* with indices in the set *G_α_*. To further resolve prey/bait interactions, we score the effect size by partitioning *G_α_* into 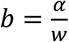 subsets {*G*_*α*1_, *G*_*α2*_,…, *G_αb_*}, where *G_αl_* = {(*i, k*): 1≤ *i* ≤ *n_b_*, 1 ≤ *k* ≤ *n_p_*, (*l* – 1)*w* ≤ *p_ik_* ≤ *lw*} for 1 ≤ *l* ≤ *b*, contains a subset of the *n_p_* * *n_b_* prey/bait combinations with similar *p*-values. The rank of the log2 fold-changes is calculated within the containing *G_αl_* subsets to obtain the fold-change enrichment score as

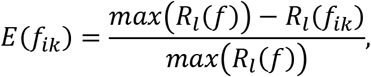

Where *R_l_*(*f*) is the rank of log2 fold-change *f_ik_* among the fold-changes with indices in set *G_αl_* and *max*(*R_l_*(*f*)) is the maximum rank of fold-changes with indices in *G_αl_*. Finally, we combined these two scores, ranking first by *p*-value and second by log2 fold-change, to obtain the enrichment score for (*i,k*) bait/prey combination as

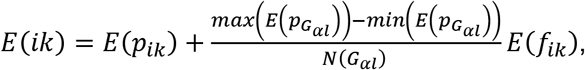

when the (*i, k*) combination is contained in *G_αl_*. Here, *E*(*p_G_αl__*) is the set of *p*-value enrichment scores with indices in *G_αl_* and *N*(*G_αl_*) is the size of *G_αl_*. Finally, to rescale *E*(*ik*) between [0,1] we divided the score by the maximum value as: 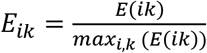.

#### Specificity score

True interactors should interact with few specific partners. To develop the specificity score we penalized preys that were enriched under selection in multiple bait screenings. Define *p_ijk_* as the *p*-value and *f_ijk_* as the log2 fold-change obtained from DESeq2, of prey *k* mated with bait *i* over prey *k* mated with bait *j* ≠ *i*, both in selected conditions. We define the *p*-value specificity score *S*(*p_ijk_*), just as *E*(*p_ik_*) was defined before. Let *G_sα_* = {(*i, j, k*): 1 ≤ *i* ≤ *n_b_, j* < *i*, 1 ≤ *k* ≤ *n_p_, p_ijk_* ≤ *α*}. Then:

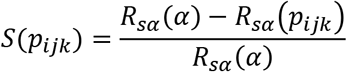

where *R_sα_*(*p*) returns the rank of *p* among the *p*-values with indices in set *G_sα_*. If *f_ijk_* < 0 or *p_ijk_* > *α* then we set *S*(*p_ijk_*) = 0. We average the *p*-value specificity scores across the *n_b_* – 1 number of bait comparisons to obtain the specificity score based in p-value of bait/prey combination (*i, k*),

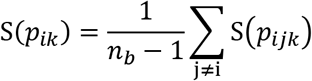

We defined *S*(*f_ijk_*), just as we did before for *E*(*f_ik_*), partitioning *G_sα_* into 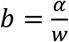 subsets {*G*_*sα*1_, *G*_*sα*2_,…, *G_sαb_*}, where *G_sαl_* = {(*i,k*): 1 ≤ *i* ≤ *n_b_, j* < *i*, 1 ≤ *k* ≤ *n_p_*, (*l* – 1)*w* ≤ *p_ijk_* ≤ *lw*} and *S*(*p_ijk_*) = 0 implying *S*(*f_ijk_*) = 0. We average over the *n_b_* – 1 number of scores *S*(*f_ijk_*) for (*i,k*) to obtain *S*(*f_ik_*):

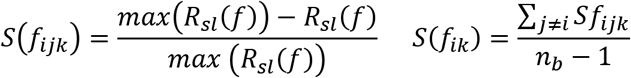

when *p_ijk_* is in *G_sαl_* and *R_sl_*(*f*) the rank of the fold-change *f* among those indexed in the *l*th subset *G_sal_*. The combined specificity score is given by:

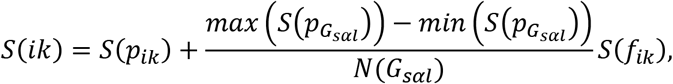

with a rescaling to [0,1], as: 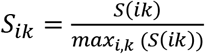.

#### In-frame score

We expect that true interactors will tend to appear *in-frame* under selective conditions. We convert the *in-frame* test of proportions statistic *ρ_ik_* into the *in-frame* score. Let *G* = {(*i, k*): 1 ≤ *i* ≤ *n_b_*, 1 ≤ *k* ≤ *n_p_*} be the complete set of prey/bait combinations. Then, the *in-frame* score is:

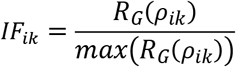

where *R_G_*(*ρ_ik_*) represents the rank of the *ρ_ik_* test statistic for prey *k* interacting with bait *i* among all prey/bait interactions. For prey with no fusion reads in either non-selected or selected conditions, *IF_ik_* was set to zero.

### Simulation of the Y2H-NGIS dataset

To test the performance of Y2H-SCORES under different conditions we developed a framework for Y2H-NGIS simulation, using empirical data to motivate the simulation model and parameter values. Fig 5 shows the experimental workflow we wish to simulate. We simulated both total and fusion read counts under selected and non-selected conditions.

**Fig 5.**
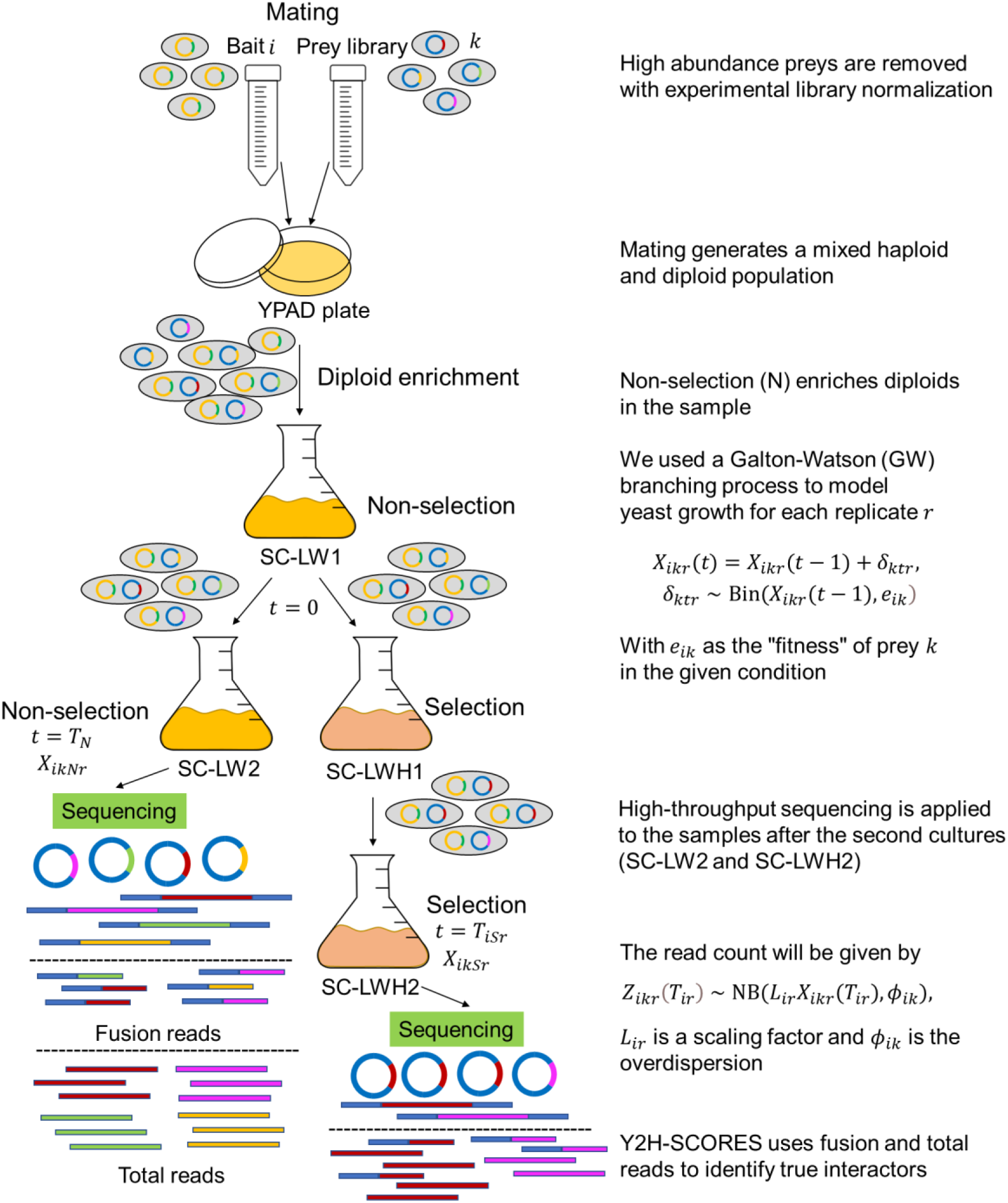
Experimental workflow for Y2H-NGIS. After the mating between bait and prey, diploids go through a non-selective culture to reach exponential phase. Once there (*t* = 0), the culture is split into two flasks, one for non-selection and another for selection. The objective of the second subculture is to grow yeast exponentially until it reaches saturation, a process that is repeated twice under selective conditions. After *T_N_* generations in the non-selected condition and *T_iSr_* generations in the selected condition, culture aliquots are taken to be sequenced.

#### Model

We used a Galton-Watson (GW) branching process to model yeast growth in each condition *c* ∈ {*S, N*}. In this presentation of the model, we drop the index *c* from the notation for simplicity. The *r^th^* replicate culture in the presence of bait *i* starts with *M_ir_*(0) = *M*_0_ = 3.84 × 10^9^ total yeast, and is grown for a potentially random number of *T_ir_* generations until the exponential growth phase ends. While the population size *M_ir_*(*T_ir_*) at the end of the experiment will be about 7.5 × 10^10^, there is enough variation in this number that we do not consider it necessary to condition on its value.

Let *X_ikr_*(*t*) be the number of yeast containing prey *k* at generation *t*. We assume *X_ikr_*(*t*) follows a simple Galton-Watson branching process,

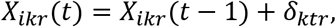

where *δ_ktr_* ~ Bin(*X_ikr_*(*t* – 1), *e_ik_*) and *e_ik_* is the “fitness” of prey *k* in the given condition with bait *i*. We will generally assume each prey is experiencing differential growth rates *e_ik_* because of selection, but the model also applies to non-selection conditions, where we assume all yeast grow at the same rate *e_ik_* = *e_N_*. The initial number of prey *k* is *X_ikr_*(0) = *M_ikr_*, with *M_ikr_* ~ Bin(*M*_0_, *q_ik_*), and given the true proportion *q_ik_* of prey *k* in the prey library.

At the end of the experiment (selection or non-selection), at generation *T_ir_*, we do not observe *X_ikr_*(*T_ir_*) directly. Instead, we observe read counts *Z_ikr_*(*T_ir_*) ~ NB(*L_ir_X_ikr_*(*T_ir_*), *ϕ_ik_*), from a Negative Binomial distribution with mean and variance:

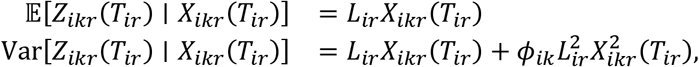

where 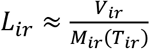 is a scaling factor (also called “size factor”) accounting for sequencing depth *V_ir_* and the population size *M_ir_*(*T_ir_*) at generation *T_ir_*. Parameter *ϕ_ik_* ≥ 0 is an overdispersion parameter that accounts for extra variation not already explained by the randomness in the initial prey count *M_ikr_* and the branching process. We treat diploid enrichment and the second round of selection as deterministic in the model (Fig 5), but they may cause overdispersion relative to our stochastic growth model. Possible overdispersion is accommodated by using an NB observation model.

For full details about the modelling and parameter estimation please refer to the S2 text file.

### Design of simulation scenarios and performance of scores

We designed different simulation scenarios to test the performance of the scores under different conditions. We took samples with 8000, 20000, and 40000 preys and 2 to 10 bait screenings. We used the following parameters to evaluate the performance scores as reported in S3 Table: **1)** the stickiness of the samples, which we define as the percentage of auto-active/non-specific preys in the library; **2)** the strength of the true interactors, defined as the minimum percentile of the observed fitness parameter 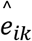 calculated from real data and used to simulate the true interactor group; **3)** overdispersion, given by the parameters *φ_kN_, φ_iks_*, and sampled randomly or from a subset with high values (over the 90^th^ percentile observed in real data); **4)** the proportion of true interactors in the prey library, given by *q_ik_*; **5)** the number of replicates; and **6)** the number of baits. S4 Fig shows sampling distributions of some of these parameters as estimated from real data. Receiver Operating Characteristic (ROC) and Precision Recall (PR) curves were constructed, and their respective Area Under the Curve (AUC) values calculated for comparison between simulations.

### Experimental procedures

We generated experimental data to estimate parameters for the simulation, and to test the efficiency of Y2H-SCORES. Using an established Gateway-compatible CEN/ARS GAL4 system (43, 44), we created a normalized, three-frame cDNA expression library of 1.1 x 10^7^ primary clones from pooled RNA isolated from a time-course experiment of barley, *Hordeum vulgare* L. (*Hv*) infected with the powdery mildew fungus, *Blumeria graminis* f. sp. *hordei* (*Bgh*) (27, 45). Baits were mated with a prey strain expressing the cDNA library and grown on selective media to identify protein-protein interactions. To initiate screening, mating of bait and prey cDNA library was performed on solid YPAD media. Diploids were enriched in SC-Leu-Trp (SC-LW) liquid media and sub-cultured under two conditions: 1) non-selected diploid growth (SC-LW) and 2) selected for reporter activation in SC-Leu-Trp-His (SC-LWH). Diploids expressing a positive PPI activate the HIS3 reporter construct and multiply in SC-LWH media whereas diploids expressing two non-interacting proteins are unable to grow under this selection. After sub-culturing the samples and reaching saturation (OD = 2.5 - 3), cells were collected, plasmids were isolated, and prey cDNA was amplified and sequenced using the Illumina HiSeq 2500 platform. We performed three independent biological replicates, collecting 5-10 million reads per sample.

Data from the *Bgh* effector protein AVR_A13_ (CSEP0372) and luciferase acting as baits were analyzed using the NGPINT pipeline. Outputs were taken to compute the Y2H-SCORES. We applied different normalization methods, calculated the Y2H-SCORES and their ensemble using Borda counts to obtain a ranked list of interactors. We compared the score values of the top 5% of the ranked interactors with each normalization method, using a Wilcoxon ranked-sum test (S4 Table). Quantiles of the specificity score for the top 100 interactions ranked using median-of-ratios normalization, and unique or non-unique across other normalizations, were calculated to show the lower values of the non-unique list (S5 Table). Using the Y2H-SCORES calculated from library size normalization and the Borda ensemble, we predicted a top list of candidate interactors to be validated. The validation consisted in identifying candidate true interactors based on the list, determining the interacting prey fragments using the IGV alignments obtained from the NGPINT pipeline (20) and the *in-frame* prey transcripts with the highest *in-frame* score. After the determination of the exact fragment, primers were designed for Gateway cloning, and subsequent insertion into the prey vector. After cloning the candidate prey into yeast, we concluded the validation with a binary Y2H test in a series of media and controls: 1) Diploid selection (SC-LW), interaction selection (SC-LWH) and stringent selection (SC-LWH+3AT) as shown in Fig 4.

### Determination of interlogs

Predicted protein-protein interactors of HvTHF1 were inferred using interlogs (32, 35). Orthologs of *HvThf1* with *Arabidopsis thaliana, Zea mays* and *Oryza sativa* were obtained using the Plant Compara tables from Ensembl Plants (46). Experimentally validated interactions for these plants were mined from BioGRID 3.5.171 version (33), the Protein-Protein Interaction database for Maize (PPIM) (47) and the Predicted Rice Interactome Network (PRIN) database (34). Barley interlogs were inferred by assigning the mined interactions from the corresponding ortholog with *Oryza sativa*, which was the only species with experimentally validated interactions reported for the THF1 protein. Visualization of the network was done using Cytoscape (48).

## Abbreviations

AUC: Area Under the Curve
*Bgh*: *Blumeria graminis* f. sp. *hordei*
CV: Coefficient of Variation
DE: Differentially Enriched
eQTL: expression Quantitative Trait Loci
*Hv*: *Hordeum vulgare*
*Mla1*: powdery mildew resistance locus a1
*MlLa*: Laevigatum resistance locus
NGPINT: **N**ext-**g**eneration **p**rotein-protein **int**eraction software
NB: Negative Binomial
PCA: Principal Component Analysis
PPI: Protein-Protein Interaction
PR: Precision Recall
ROC: Receiver Operating Characteristic
ROS: Reactive oxygen species
RUV: Remove Unwanted Variation
TPM: Transcripts Per Million
Y2H: Yeast Two-Hybrid
Y2H-NGIS: Yeast Two-Hybrid Next-Generation Interaction Screening

## Availability of code, data, and materials

R code and ReadMe file for the Y2H-SCORES software are provided at GitHub (https://github.com/Wiselab2/Y2H-SCORES). Additionally, we implemented a python script to link the score functions with the NGPINT pipeline (20), which integrates both software packages by allowing Y2H-SCORES to run on the NGPINT outputs. Users can find the instructions in the same repository. Y2H-NGIS data supporting the conclusions of this article will be available in NCBI’s Gene Expression Omnibus (GEO) upon publication.

## Supplemental files

**S1 Fig.** Coefficient of variation (CV) for each prey using different normalization methods.

**S2 Fig.** Top candidate interactors inferred with Y2H-SCORES and different normalization methods.

**S3 Fig.** Binary Y2H for the candidate preys HORVU2Hr1G060120 and HORVU2Hr1G024160.

**S4 Fig.** Distributions of the parameters used for Y2H-NGIS simulation.

**S1 Table.** Wilcoxon signed-rank test for the Pearson correlation of samples grouped by condition, for different normalization methods.

**S2 Table.** Wilcoxon signed-rank test for CV densities of different normalization methods.

**S3 Table.** Y2H-NGIS simulation scenarios and performance of Y2H-SCORES.

**S4 Table.** Wilcoxon ranked-sum test for the top 5% ranked interactions using the Borda ensemble of the Y2H-SCORES under different normalization methods.

**S5 Table.** Quantiles of the specificity score for the top 100 interactions ranked using median-of-ratios normalization and unique or non-unique across other normalizations.

**S1 Text.** *HvThf1* prey sequences.

**S2 Text.** Y2H-NGIS simulation model.

## Acknowledgments

The authors thank Greg Fuerst for conducting the time-course infection experiment and for <u>expert</u> isolation of RNA for the 3-frame Y2H library. Research supported in part by Fulbright - Minciencias 2015 & Schlumberger Faculty for the Future fellowships to VVZ, USDA-NIFA-ELI Postdoctoral Fellowship 2017-67012-26086 to JME, Oak Ridge Institute for Science and Education (ORISE) under U.S. Department of Energy (DOE) contract number DE-SC0014664 to SB, USDA-National Institute of Food and Agriculture (NIFA) Hatch project IOW03617 to KSD, and National Science Foundation - Plant Genome Research Program grant 13-39348 and USDA-Agricultural Research Service project 3625-21000-067-00D to RPW. The funders had no role in study design, data collection and analysis, decision to publish, or preparation of the manuscript. Mention of trade names or commercial products in this publication is solely for the purpose of providing specific information and does not imply recommendation or endorsement by the USDA, NIFA, ARS, DOE, ORAU/ORISE or the National Science Foundation. USDA is an equal opportunity provider and employer.

