## Supplementary material for "Y2H-SCORES: A statistical framework to infer protein-protein interactions from next-generation yeast-two-hybrid sequence data": S2 Figure

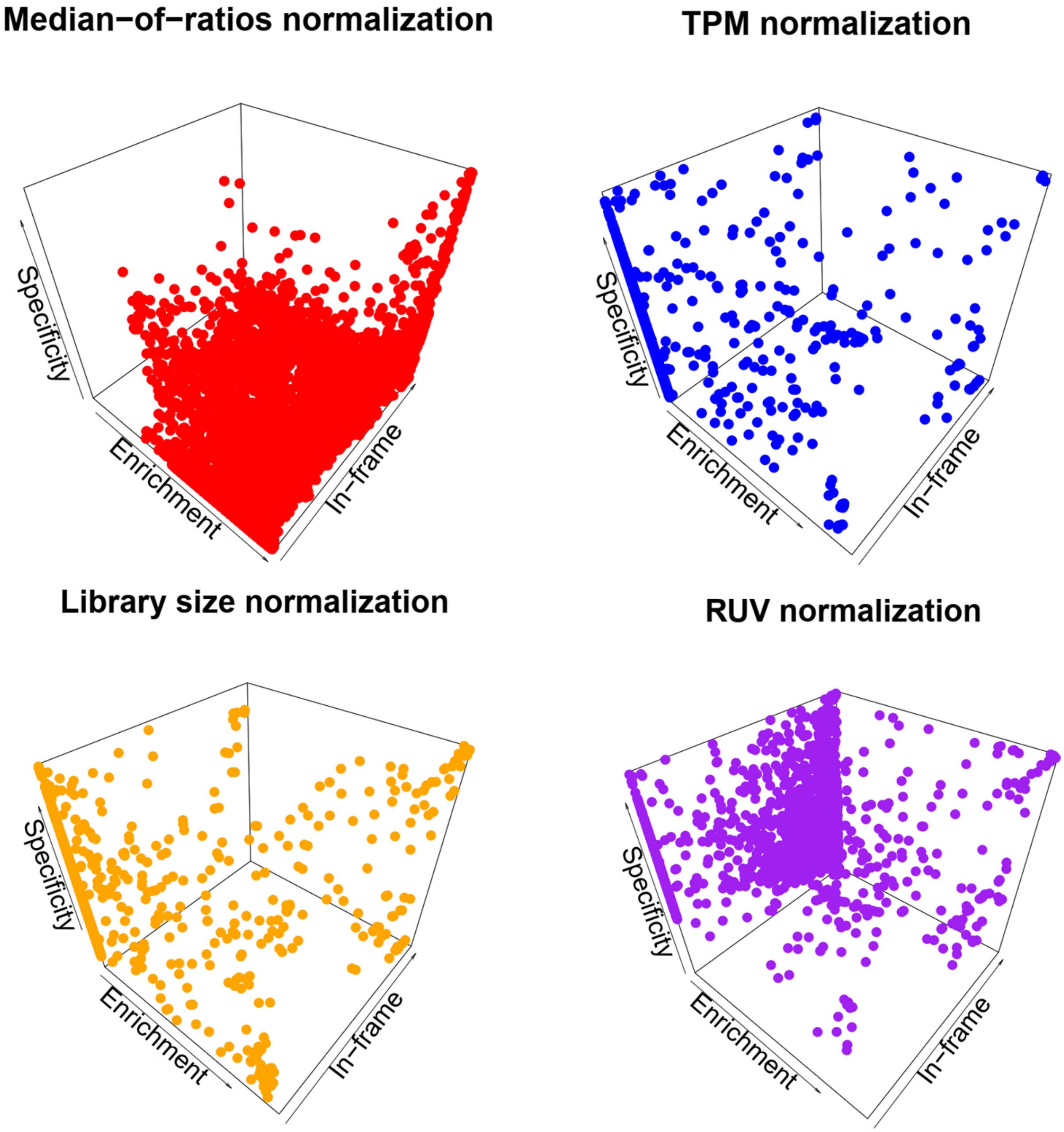


S2 Fig. Top candidate interactors inferred with Y2H-SCORES and different normalization methods. Only top 5% Borda score values are shown.

Velásquez-Zapata et. al. (2020) Y2H-SCORES: A statistical framework to infer protein-protein interactions from next-generation yeast-two-hybrid sequence data.
