## Supplementary material for "Y2H-SCORES: A statistical framework to infer protein-protein interactions from next-generation yeast-two-hybrid sequence data": S3 Figure

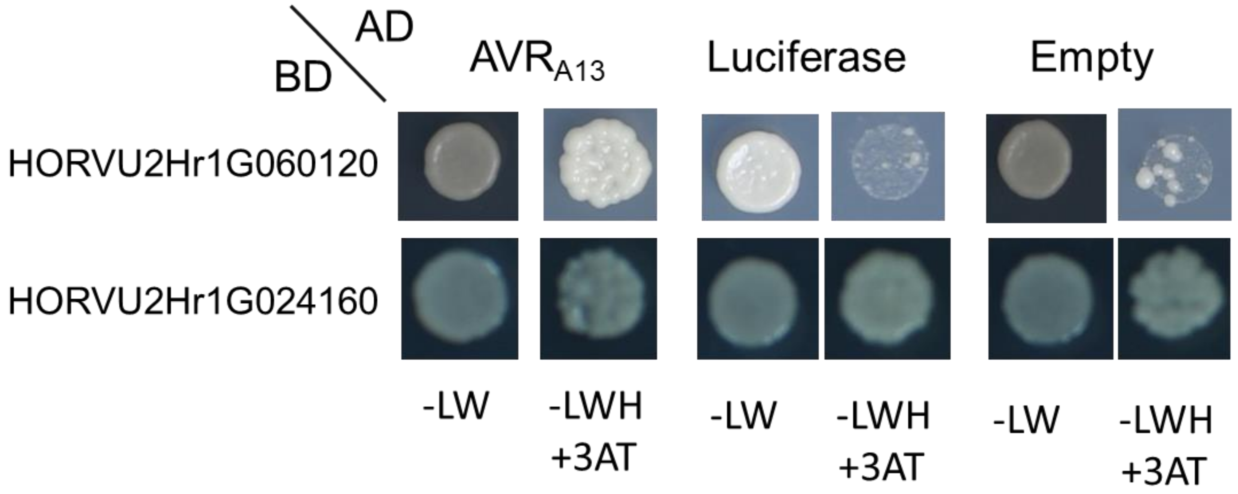


S3 Fig. Binary Y2H for the candidate preys HORVU2Hr1G060120 and HORVU2Hr1G024160. SC-LW media was used as control for diploid growth, and the interaction was tested using stringent selection with 3-amino-1,2,4-triazole (3AT) in SC-LWH media. Positive interaction tests with AVR$_{A13}$, luciferase and empty bait confirm these preys are auto-active.

Velásquez-Zapata et. al. (2020) Y2H-SCORES: A statistical framework to infer protein-protein interactions from next-generation yeast-two-hybrid sequence data.
