## Supplementary material for "Y2H-SCORES: A statistical framework to infer protein-protein interactions from next-generation yeast-two-hybrid sequence data": S1 Figure

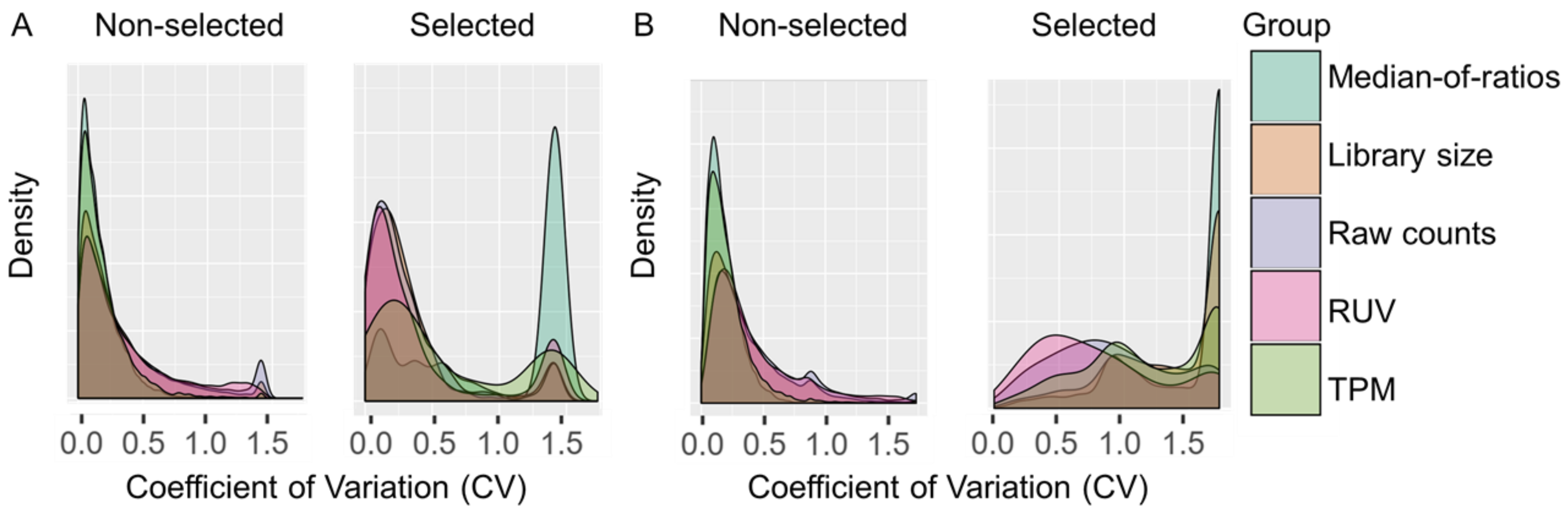


S1 Fig. Coefficient of variation (CV) for each prey using different normalization methods (color coded). Higher CV values may indicate poor performance because of a high variation between replicates. Baits: A) Luciferase and B) MLA6-LRR.

Velásquez-Zapata et. al. (2020) Y2H-SCORES: A statistical framework to infer protein-protein interactions from next-generation yeast-two-hybrid sequence data.
