## Supplementary material for "Y2H-SCORES: A statistical framework to infer protein-protein interactions from next-generation yeast-two-hybrid sequence data": S4 Figure

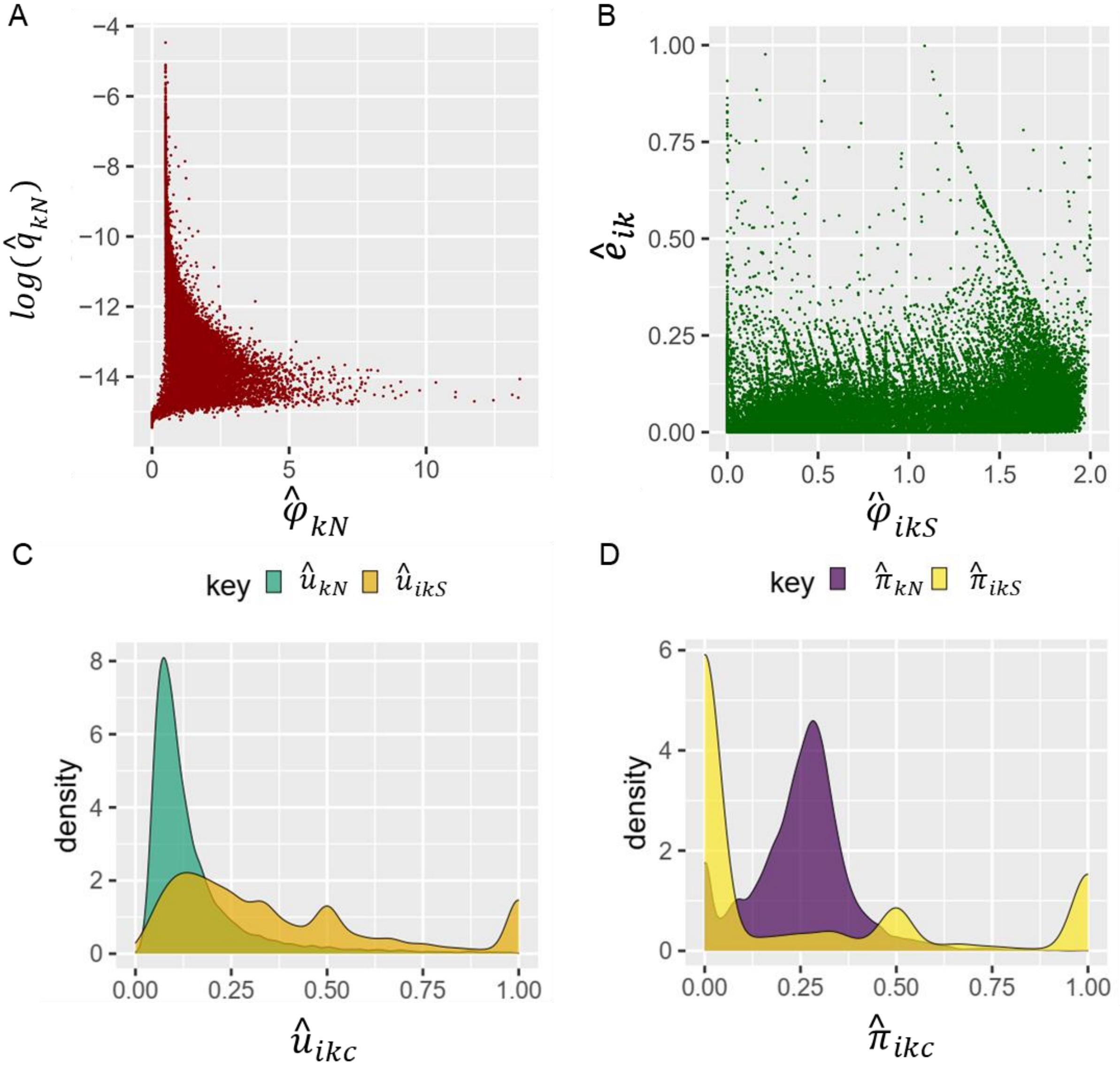


S4 Fig. Distributions of the parameters used for Y2H-NGIS simulation. A) The prey proportions and the overdispersion in non-selected samples. B) Fitness coefficient and overdispersion in selected samples. C) The proportion of fusion reads in the samples. D) the proportion of *in-frame* reads in the samples.

Velásquez-Zapata et. al. (2020) Y2H-SCORES: A statistical framework to infer protein-protein interactions from next-generation yeast-two-hybrid sequence data.
