## Supplementary material for "Y2H-SCORES: A statistical framework to infer protein-protein interactions from next-generation yeast-two-hybrid sequence data": S2 Text

### S2 Text. Y2H-NGIS Simulation.

#### Simulation

To test the performance of Y2H-SCORES under different conditions we developed a framework for Y2H-NGIS simulation, using empirical data to motivate simulation models and parameter values. Figure 5 shows the experimental workflow we wish to simulate. We simulated both total and fusion read counts under selected and non-selected conditions.

In what follows, we will explain how non-selected, selected, and in-frame prey counts were simulated. We explain how every parameter of our model is estimated from real data. During the simulation study, the most important variables will be intentionally varied to assess their impact. Key to these simulations is a model for exponential yeast growth, which we describe next.

##### Model

We used a Galton-Watson (GW) branching process to model yeast growth in each condition $c \in \{S, N\}$. In this presentation of the model, we drop the index $c$ from the notation for simplicity. The $r$th replicate culture in the presence of bait $i$ starts with $M_{ir}(0) = M_0 = 3.84 \times 10^9$ total yeast, and is grown for a potentially random number of $T_{ir}$ generations until the exponential growth phase ends. While the population size $M_{ir}(T_{ir})$ at the end of the experiment will be about $7.5 \times 10^{10}$, there is enough variation in this number that we do not consider it necessary to condition on its value.

Let $X_{ikr}(t)$ be the number of yeast containing prey $k$ at generation $t$. We assume $X_{ikr}(t)$ follows a simple Galton-Watson branching process,

$$X_{ikr}(t) = X_{ikr}(t-1) + \delta_{ktr},$$

where $\delta_{ktr} \sim \text{Bin}(X_{ikr}(t-1), e_{ik})$ and $e_{ik}$ is the "fitness" of prey $k$ in the given condition with bait $i$. We will generally assume each prey is experiencing differential growth rates $e_{ik}$ because of selection, but the model also applies to non-selection conditions, where we assume all yeast grow at the same rate $e_{ik} = e_N$.

From branching process theory, we know

$$\begin{aligned}\mathbb{E}\left[X_{ikr}(t) \mid X_{ikr}(0) = 1\right] &= (1 + e_{ik})^t \\ \text{Var}\left[X_{ikr}(t) \mid X_{ikr}(0) = 1\right] &= (1 - e_{ik})(1 + e_{ik})^{t-1}\left[(1 + e_{ik})^t - 1\right].\end{aligned}$$

By the independence of yeast during exponential growth, if the initial number of prey $k$ is $X_{ikr}(0) = M_{ikr}$, then

$$\begin{aligned}\mathbb{E}\left[X_{ikr}(t) \mid X_{ikr}(0) = M_{ikr}\right] &= M_{ikr}(1 + e_{ik})^t \\ \text{Var}\left[X_{ikr}(t) \mid X_{ikr}(0) = M_{ikr}\right] &= M_{ikr}(1 - e_{ik})(1 + e_{ik})^{t-1}\left[(1 + e_{ik})^t - 1\right].\end{aligned}$$

Given the true proportion $q_{ik}$ of prey $k$ in the prey library, the initial number of prey $k$ yeast,

$$M_{ikr} \sim \text{Bin}(M_0, q_{ik}),$$

is a binomial random variable, where $M_0$ is the initial number of diploid yeast cells starting the culture. By the Laws of Total Expectation and Total Variance, we have the unconditional expectation and variance are

$$\begin{aligned}\mathbb{E}\left[X_{ikr}(t)\right] &= M_0 q_{ik}\left(1+e_{ik}\right)^t \\ \mathrm{Var}\left[X_{ikr}(t)\right] &= M_0 q_{ik}\left(1-e_{ik}\right)\left(1+e_{ik}\right)^{t-1}\left[\left(1+e_{ik}\right)^t-1\right]+\left(1+e_{ik}\right)^{2t} M_0 q_{ik}\left[1-q_{ik}\right].\end{aligned}$$

At the end of the experiment (selection or non-selection), at generation $T_{ir}$, we do not observe $X_{ikr}(T_{ir})$ directly. Instead, we observe read counts

$$Z_{ikr}\left(T_{ir}\right) \sim \mathrm{NB}(L_{ir} X_{ikr}(T_{ir}), \phi_{ik}),$$

from a Negative Binomial distribution with mean and variance

$$\begin{aligned}\mathbb{E}\left[Z_{ikr}\left(T_{ir}\right) \mid X_{ikr}\left(T_{ir}\right)\right] &= L_{ir} X_{ikr}\left(T_{ir}\right) \\ \mathrm{Var}\left[Z_{ikr}\left(T_{ir}\right) \mid X_{ikr}\left(T_{ir}\right)\right] &= L_{ir} X_{ikr}\left(T_{ir}\right)+\phi_{ik} L_{ir}^2 X_{ikr}^2\left(T_{ir}\right)\end{aligned}$$

where $L_{ir} \approx \frac{V_{ir}}{M_{ir}(T_{ir})}$ is a scaling factor (also called "size factor") accounting for sequencing depth $V_{ir}$ and the population size $M_{ir}(T_{ir})$ at generation $T_{ir}$. Parameter $\phi_{ik} \geq 0$ is an overdispersion parameter that accounts for extra variation not already explained by the randomness in the initial prey count $M_{ikr}$ and the branching process. We treat diploid enrichment and the second round of selection as deterministic in the model (Fig 5), but either may cause overdispersion relative to our stochastic growth model. Possible overdispersion is accommodated by using a NB observation model.

The assumption $L_{ir} = \frac{V_{ir}}{M_{ir}(T_{ir})}$ is good if PCR amplification plays a negligible role in the sampling, meaning read depth is much smaller than the number of yeast cells sampled and there is no bias in the amplification. The unconditional mean and variance are therefore

$$\begin{aligned}\mathbb{E}\left[Z_{ikr}\left(T_{ir}\right)\right] &= L_{ir}\,\mathbb{E}\left[X_{ikr}\left(T_{ir}\right)\right] \\ \mathrm{Var}\left[Z_{ikr}\left(T_{ir}\right)\right] &= L_{ir}\,\mathbb{E}\left[X_{ikr}\left(T_{ir}\right)\right]+\phi_{ik} L_{ir}^2\,\mathbb{E}\left[X_{ikr}^2\left(T_{ir}\right)\right]+L_{ir}^2 \mathrm{Var}\left[X_{ikr}\left(T_{ir}\right)\right].\end{aligned}$$

#### Estimation of parameters

The proportion of true interactors in the library varies with bait, but is roughly $\mathrm{Unif}(0.0004, 0.001)$. This distribution is based on the data of Pashova et al. (2016) who confirmed 8 out of ~15000 preys to be true interactors. Our experiments also showed a similar trend, having confirmed interactions between 1 and 25 in a ~36000 prey population.

Observed size factors were computed as $\hat{L}_{icr} = \frac{v_{icr}}{M_{icr}(T_{icr})}$ for the $r$th replicate of bait $i$ under condition $c$. Here, $v_{icr} = \sum_{k=1}^{K} z_{ikcr}(T_{icr})$ is the observed coverage and $M_{icr}(T_{icr})$ is the yeast population size at the end of the experiment. For this calculation, we assumed the rough estimate $M_{icr}(T_{icr}) = 7.5 \times 10^{10}$, which is constant across baits, conditions and replicates.

##### No selection

Under no selection, each prey should replicate at the same rate regardless of bait, and we can drop the bait index $i$. Furthermore, we assume $e_N = 1$ under these ideal conditions, so every yeast replicates at every generation. Because the initial population size $M_{iNr}(0) = M_0 = 3.84 \times 10^9$ is large and all yeast are actively replicating, the total population size $M_{iNr}(t) = M_N(t)$ is not only independent of bait $i$, but also effectively deterministic and thus independent of replicate $r$, even if the individual number $M_{ikNr}(t)$ of some prey $k$ is notably stochastic and not constant with bait $i$ and replicate $r$. Since the experiment is stopped based on the total population size, the number of elapsed generations $T_{iNr} = T_N$ is also constant across baits and replicates in the non-selection experiments. Under the branching process formulation, after $t$ generations, the population size is expected to reach

$$\mathbb{E}\left[M_N(t) \mid M_0\right] = M_0(1+e_N)^t.$$

Given information about the initial population size $M_0$ and the final population size $M_N(T_N)$, we can estimate the number of elapsed generations as

$$T_N = \log_2 \left( \frac{M_N(T_N)}{M_0} \right) = \log_2 \left( \frac{7.5 \times 10^{10}}{3.84 \times 10^9} \right) \approx 4.29,$$

which we round to $T_N = 4$ in practice.

Furthermore, we expect the proportion of prey $k$ in the population $q_{ikNr}(t)$ at generation $t$ to satisfy

$$\mathbb{E}\left[q_{ikNr}(t) \mid q_{ikNr}(0)\right] = q_{ikNr}(0),$$

and the expected initial proportion $\mathbb{E}\left[q_{ikNr}(0)\right] = q_k$ is determined by the prey library, independent of replicate $r$, bait $i$ and experimental condition ($S$ or $N$). We have observed $z_{ikNr}(T_N)$ reads of prey $k$ in the presence of the $r$th replicate of bait $i$ in the absence of selection. We can use the method of moments to estimate

$$\hat{q}_k = \frac{1}{\sum_{i,r} 1} \sum_{i,r} \frac{z_{ikNr}\left(T_N\right)}{v_{iNr}},$$

the average proportion of prey $k$ observed across all baits and replicates in the non-selection condition.

We can also use the method of moments to estimate overdispersion parameters $\phi_{kN}$ for each prey. Estimating equation

$$\begin{aligned}
\sum_{i,r} z^2_{ikNr}(T_N) &= \sum_{i,r} \mathrm{Var}(Z_{ikNr}(T_N)) + \mathbb{E}[Z_{ikNr}(T_N)]^2 \\
&= \sum_{i,r} L_{iNr}\, \mathbb{E}\left[X_{ikNr}\left(T_N\right)\right] + \hat{\phi}_{kN} L^2_{iNr}\, \mathbb{E}\left[X^2_{ikNr}\left(T_N\right)\right] \\
&\quad + L^2_{iNr} \mathrm{Var}\left[X_{ikNr}\left(T_N\right)\right] + \left(L_{iNr}\, \mathbb{E}\left[X_{ikNr}\left(T_N\right)\right]\right)^2 \\
&= \sum_{i,r} L_{iNr}\, \mathbb{E}\left[X_{ikNr}\left(T_N\right)\right] + \hat{\phi}_{kN} L^2_{iNr} (\mathrm{Var}\left[X_{ikNr}\left(T_N\right)\right] + \mathbb{E}\left[X_{ikNr}\left(T_N\right)\right]^2) \\
&\quad + L^2_{iNr} \mathrm{Var}\left[X_{ikNr}\left(T_N\right)\right] + \left(L_{iNr}\, \mathbb{E}\left[X_{ikNr}\left(T_N\right)\right]\right)^2,
\end{aligned}$$

yields estimate

$$\hat{\phi}_{kN} = \frac{\sum_{i,r} \left\{ z^2_{ikNr}(T_N) - L^2_{iNr} \mathrm{Var}[X_{ikNr}\left(T_N\right)] - L_{iNr}\, \mathbb{E}[X_{ikNr}\left(T_N\right)] - \left(L_{iNr}\, \mathbb{E}[X_{ikNr}\left(T_N\right)]\right)^2 \right\}}{\sum_{i,r} \left\{ L^2_{iNr} \left( \mathrm{Var}[X_{ikNr}\left(T_N\right)] + \mathbb{E}[X_{ikNr}\left(T_N\right)]^2 \right) \right\}},$$

with

$$\begin{aligned}
\mathbb{E}\left[X_{ikNr}(T_N)\right] &= M_0 \hat{q}_k \left(1 + e_N\right)^{T_N} \\
\mathrm{Var}\left[X_{ikNr}(T_N)\right] &= M_0 \hat{q}_k \left(1 - e_N\right) \left(1 + e_N\right)^{T_N - 1} \left[\left(1 + e_N\right)^{T_N} - 1\right] \\
&\quad + \left(1 + e_N\right)^{2T_N} M_0 \hat{q}_k \left[1 - \hat{q}_k\right].
\end{aligned}$$

##### Selection

By experimental design, we know the initial number of cells $M_{iSr}(0) = M_0 = 3.84 \times 10^9$ in the selection experiments, which do not vary by bait $i$ or replicate $r$. Let $q_{ikSr}(0)$ be the initial proportion of prey $k$ at generation $t = 0$ in the selection experiment with bait $i$. Since both non-selected and selected growth were initialized in the same way and non-selected growth does not change the prey proportions, we can assume the initial prey proportion $q_{ikSr}(0) = \hat{q}_k$, where $\hat{q}_k$ were estimated from the no selection experiments. Then, the initial number of prey $k$ yeast in the selection replicate $r$ against bait $i$,

$$M_{ikSr}(0) \sim \mathrm{Bin}\left(M_0, \hat{q}_k\right),$$

is a binomial random variable.

Culture growth under selection depends on the bait and because it is so severely inhibited, becomes stochastic, so the time to saturation is observed to vary across baits and replicates. We estimated the number of generations $T_{iSr} = h_{iSr}/g_T$ per selection experiment assuming a constant generation time $g_T = h_N/T_N$, calculated from the observed culture time $h_N = 18h$ of the non-selected samples, which was constant across experiments, and the observed culture times $h_{iSr}$ of selected samples, varying across baits and replicates.

By the end of the experiment, after $T_{iSr}$ generations of growth, the theory developed earlier yields unconditional mean and variance

$$\begin{aligned}\mathbb{E}\left[Z_{ikSr}\left(T_{iSr}\right)\right] &= L_{iSr}\,\mathbb{E}\left[X_{ikSr}\left(T_{iSr}\right)\right] \\ \mathrm{Var}\left[Z_{ikSr}\left(T_{iSr}\right)\right] &= L_{iSr}\,\mathbb{E}\left[X_{ikSr}\left(T_{iSr}\right)\right] + \phi_{ikS}L_{iSr}^2\,\mathbb{E}\left[X_{ikSr}^2\left(T_{iSr}\right)\right] + L_{iSr}^2\mathrm{Var}\left[X_{ikSr}\left(T_{iSr}\right)\right]\end{aligned}$$

for the observed count $Z_{ikSr}\left(T_{iSr}\right)$ of prey $k$.

We used the method of moments to estimate $e_{ik}$ from the observed counts $z_{ikSr}(T_{iSr})$ as

$$\sum_{r=1}^{3} z_{ikSr}(T_{iSr}) = \sum_{r=1}^{3} L_{iSr}M_0\hat{q}_k\left(1+\hat{e}'_{ik}\right)^{T_{iSr}},$$

which yields $\hat{e}'_{ik}$ as the root of equation

$$0 = \sum_{r=1}^{3} L_{iSr}M_0\hat{q}_k\left(1+\hat{e}'_{ik}\right)^{T_{iSr}} - \sum_{r=1}^{3} z_{ikSr}(T_{iSr}),$$

equivalent to

$$0 = \frac{M_0}{M_{iSr}(T_{iSr})}\sum_{r=1}^{3} \hat{q}_k\left(1+\hat{e}'_{ik}\right)^{T_{iSr}} - \sum_{r=1}^{3} \hat{q}_{ikSr}(T_{iSr}),$$

where once again we make the assumption that $M_{iSr}(T_{iSr}) \approx 7.5 \times 10^{10}$ at the end of the experiment, and thus is constant across baits and replicates. We then estimated $\hat{e}'_{ik}$ using the Newton-Raphson method implemented by the uniroot function in R. To assure a positive estimate, we ultimately set

$$\hat{e}_{ik} = \max\left\{0, \hat{e}'_{ik}\right\}.$$

As in the non-selection experiments, we can estimate the overdispersion parameters $\phi_{ikS}$ as

$$\hat{\phi}_{ikS} = \frac{\sum_{r=1}^{3}\left\{z_{ikSr}^2(T_{iSr}) - L_{iSr}^2\mathrm{Var}[X_{ikSr}\left(T_{iSr}\right)] - L_{iSr}\,\mathbb{E}[X_{ikSr}\left(T_{iSr}\right)] - \left(L_{iSr}\,\mathbb{E}[X_{ikSr}\left(T_{iSr}\right)]\right)^2\right\}}{\sum_{r=1}^{3}\left\{L_{iSr}^2\left(\mathrm{Var}[X_{ikSr}\left(T_{iSr}\right)] + \mathbb{E}[X_{ikSr}\left(T_{iSr}\right)]^2\right)\right\}}.$$

with

$$\begin{aligned}\mathbb{E}\left[X_{ikSr}(T_{iSr})\right] &= M_0\hat{q}_k\left(1+\hat{e}_{ik}\right)^{T_{iSr}} \\ \mathrm{Var}\left[X_{ikSr}(T_{iSr})\right] &= M_0\hat{q}_k\left(1-\hat{e}_{ik}\right)\left(1+\hat{e}_{ik}\right)^{T_{iSr}-1}\left[\left(1+\hat{e}_{ik}\right)^{T_{iSr}} - 1\right] \\ &\quad + \left(1+\hat{e}_{ik}\right)^{2T_{iSr}} M_0\hat{q}_k\left[1-\hat{q}_k\right].\end{aligned}$$

#### Fusion data

We expect some fraction $u_{ikc}$ of the $Z_{ikcr}(T_{icr})$ observed reads to be fusion reads $F_{ikcr}$. The factors determining $u_{ikc}$ include the placement of the PCR primers, the read length and coverage, the library and bait in selective conditions. It is difficult to model this process, but in a high throughput experiment, there are enough prey to model the distribution empirically. After verifying that the fusion read fraction was stable across baits and replicates in non-selection conditions, we simply estimated $\hat{u}_{kN} = \frac{1}{\sum_{i,r} 1}\sum_{i,r}\frac{f_{ikNr}}{z_{ikNr}(T_N)}$.

In selected conditions, the fusion read fraction was stable across replicates, but not baits, so we estimated $\hat{u}_{ikS} = \frac{1}{\sum_r 1} \sum_r \frac{f_{ikSr}}{z_{ikSr}(T_{iSr})}$. The resulting empirical distributions are shown in Figure S4.

If the number of *in-frame* fusion reads is given by $Y_{ikcr}$, some proportion $\pi_{ikc} = \frac{Y_{ikcr}}{F_{ikcr}}$ of the fusion reads will be *in-frame*. Again, we observed little variation across baits and replicates in non-selection conditions and little variation across replicates in selection conditions. For the non-selected condition, we computed the observed proportions $\hat{\pi}_{kN} = \frac{1}{\sum_{i,r} 1} \sum_{i,r} \frac{y_{ikNr}}{f_{ikNr}}$ for all prey with at least one fusion read. We used the density $\hat{\pi}_{kN}$ to sample $\pi_{kN}$ during the simulation. Similarly, for the selected condition we calculated $\hat{\pi}_{ikS} = \frac{1}{\sum_r 1} \sum_r \frac{y_{ikSr}}{f_{ikSr}}$ and used it to sample $\pi_{ikS}$.

For selection conditions, the *in-frame* proportion $\pi_{ikS}$ will further depend on the bait and the type of prey, true interactor, non-interactor, or auto-active/non-specific interactor. Since we do not know the true interactor status of each prey, we set true interactors $\pi_{ikS} \sim \hat{\pi}_{ikS} \; \{\pi_{ikS} \in \hat{\pi}_{ikS} : \hat{\pi}_{ikS|0.95} < \pi_{ikS} < \hat{\pi}_{ikS}\}$, which matches the right peak of the observed *in-frame* proportions in selected conditions (Fig 2 and Fig S4D). In the case of non-interactors and auto-active/non-specific interactors, we sampled from this density without restiction. S4 Fig shows these distributions depicting a frame preference during selection, which points to a higher efficiency of a specific prey frame during selection. In the case of true interactors we asume this frame should be *in-frame* to have a biological meaning.

#### Simulation algorithm

Our simulation relies on parameters estimated from real data, namely estimates

$$
\begin{aligned}
\mathcal{Q}_N &= \left\{(\hat{q}_k, \hat{\phi}_{kN}) : 1 \le k \le K\right\} \text{ from the non-selection experiments} \\
\mathcal{Q}_S &= \left\{(\hat{e}_{ik}, \hat{\phi}_{ikS}) : 1 \le i \le I, 1 \le k \le K\right\} \text{ from the selection experiments, and} \\
\mathcal{L} &= \left\{\hat{L}_{icr} : 1 \le i \le I, 1 \le r \le 3, C \in \{S, N\}\right\} \text{ from all experiments} \\
\mathcal{T}_S &= \left\{\hat{T}_{iSr} : 1 \le i \le i, 1 \le r \le 3\right\} \text{ from all selection experiments} \\
\mathcal{U}_N &= \{\hat{u}_{kN} : 1 \le k \le K\} \text{ from all non-selection experiments} \\
\mathcal{U}_S &= \{\hat{u}_{ikS} : 1 \le i \le I, 1 \le k \le K\} \text{ from all selection experiments.} \\
\mathcal{P}_N &= \{\hat{\pi}_{kN} : 1 \le k \le K\} \text{ from all non-selection experiments} \\
\mathcal{P}_S &= \{\hat{\pi}_{ikS} : 1 \le i \le I, 1 \le k \le K\} \text{ from all selection experiments.}
\end{aligned}
$$

In addition, the simulation takes several parameters as input: the proportion of auto-active/non-specific prey $p_s$, the fitness threshold $e_t$, the number of simulated baits $I_{sim}$, the total number of prey $n_p$, the initial selected population size and the initial non-selected population size $M_{iSr}(0) = M_N(0) = M_0$, and the final population size $M_N(T_N)$, assumed to be approximately equal to the final selected population sizes $M_{iSr}(T_{iSr})$. The number $T_N$ of generations elapsed in non-selected conditions is implied by these choices. True interactors are assumed to have fitness $e_{ik} \ge e_t$, while non-interactors have $e_{ik} < e_t$. A proportion $p_s$ of auto-active/non-specific interactors in the sample, which autoactivate in the presence of any bait, or interact with multiple baits. Their $e_{ik}$ are selected from the top 10% of $\hat{e}_{ik} < e_t$ in $\mathcal{Q}_S$.

Select $n_i \sim \text{Unif}(0.0004, 0.001) \cdot n_p$ true interactors and $n_s \sim p_s(n_p - n_i)$ auto-active/non-specific interactors from the non-interactors. Compute the 99th percentile $e_{0.99t}$ of all fitness parameters $\hat{e}_{ik} < e_t$. Compute $T_N = \log_2\left(\frac{M_N(T_N)}{M_0}\right)$ In what follows, $\text{GW}(M, e, t)$ is the Galton-Walton random branching process initialized with $M$ particles, replicating at rate $e$ for $t$ generations. For the fusion count simulation we calculate the 95th percentile $\pi_{ikS|0.95}$ of all the *in-frame* proportion for selected condition $\pi_{ikS|0.95} < \hat{\pi}_{ikS}$.

For each bait $1 \le j \le I_{\text{sim}}$:

- For each replicate $r$,
  - select $L_{jSr}, L_{jNr}$ from $\mathcal{L}$ with replacement, and

  - select $T_{jSr}$ from $\mathcal{T}_S$ with replacement.
- For each prey $l$ in the non-selected condition:
  - If $j = 1$, this is the first prey, we select parameters that will be shared by subsequent preys:
    * Select $(q_l, \phi_{lN})$ from subset $\mathcal{Q}_{\mathcal{N}}$ with replacement unless we want to simulation low abundance interactors, in which case truncate $q_l$ above the minumum values sampled during the simulation $\sim 1 \times 10^{-8}$ if $l$ is a true interactor, with $\phi_{lN} = 0$.
    * If we want to simulate high overdispersion then we sample $\phi_{lN}$ from $\left\{\hat{\phi}_{kN} \in \mathcal{Q}_{\mathcal{N}} : \hat{\phi}_{kN|0.9} < \hat{\phi}_{kN}\right\} \subset \mathcal{Q}_N$
    * Sample proportion of fusion reads $u_{lN}$ from $\mathcal{U}_N$.
    * Sample proportion of *in-frame* reads $\pi_{lN}$ from $\mathcal{P}_N$.
  - For each replicate $r$:
    * Simulate $X_{jlNr}(T_N) \sim \mathrm{GW}\left(M_0 q_l, e_N, T_N\right)$.
    * Simulate $Z_{jlNr}(T_N) \sim \mathrm{NB}\left(L_{jNr} X_{jlNr}(T_N), \phi_{lN}\right)$.
    * Simulate number of fusion reads $F_{jlNr} \sim \mathrm{Bin}\left(Z_{jlNr}(T_N), u_{lN}\right)$.
    * Simulate number of *in-frame* fusion reads $Y_{jlNr} \sim \mathrm{Bin}\left(F_{jlNr}, \pi_{lN}\right)$.
- For each prey $l$ in the selected condition:
  - Sample proportion of fusion reads $u_{jlS}$ from $\mathcal{U}_S$.
  - If $l$ is true interactor with bait $j$,
    * Select $(e_{jl}, \phi_{jlS})$ from subset $\left\{\left(\hat{e}_{ik}, \hat{\phi}_{ikS}\right) \in \mathcal{Q}_{\mathcal{S}} : \hat{e}_{ik} > e_t\right\} \subset \mathcal{Q}_S$.
    * Sample $\pi_{jlS}$ from subset $\left\{\hat{\pi}_{ikS}, \in \mathcal{P}_S : \hat{\pi}_{ikS} > \hat{\pi}_{ikS|0.95}\right\} \subset \mathcal{P}_S$.
  - Else if it is a auto-active/non-specific interactor
    * Select $(e_{jl}, \phi_{jlS})$ from subset $\left\{\left(\hat{e}_{ik}, \hat{\phi}_{ikS}\right) \in \mathcal{Q}_{\mathcal{S}} : \hat{e}_{0.99t} < \hat{e}_{ik} < e_t\right\} \subset \mathcal{Q}_S$.
    * Sample $\pi_{jlS}$ from $\mathcal{P}_S$.
  - Else if it is a non-interactor and $j = 1$,
    * Select $(e_{jl}, \phi_{jlS})$ from subset $\left\{\left(\hat{e}_{ik}, \hat{\phi}_{ikS}\right) \in \mathcal{Q}_{\mathcal{S}} : \hat{e}_{ik} \leq e_t\right\} \subset \mathcal{Q}_S$ and set $k'$ to the prey $k$ selected.
    * Sample $\pi_{jlS}$ from $\mathcal{P}_S$.
  - Else if it is a non-interactor or a true interactor and $j > 1$,
    * Select $(e_{jl}, \phi_{jlS})$ from subset $\left\{\left(\hat{e}_{ik}, \hat{\phi}_{ikS}\right) \in \mathcal{Q}_{\mathcal{S}} : k = k'\right\}$.
    * Sample $\pi_{jlS}$ from $\mathcal{P}_S$.
  - If we want to simulate high overdispersion then we sample $\phi_{jlS}$ from $\left\{\hat{\phi}_{ikS} \in \mathcal{Q}_{\mathcal{S}} : \hat{\phi}_{ikS|0.9} < \hat{\phi}_{ikS}\right\} \subset \mathcal{Q}_S$
  - For each replicate $r$:
    * Simulate $X_{jlSr}(T_{jSr}) \sim \mathrm{GW}\left(M_0 q_l, e_{jl}, T_{jSr}\right)$.
    * Simulate $Z_{jlSr}(T_{jSr}) \sim \mathrm{NB}\left(L_{jSr} X_{jlSr}(T_{jSr}), \phi_{jlS}\right)$.
    * Simulate number of fusion reads $F_{jlSr} \sim \mathrm{Bin}\left(Z_{jlSr}(T_{jSr}), u_{jlS}\right)$.
    * Simulate $Y_{jlSr} \sim \mathrm{Bin}\left(F_{jlSr}, \pi_{jlS}\right)$.
