## Supplementary material for "Y2H-SCORES: A statistical framework to infer protein-protein interactions from next-generation yeast-two-hybrid sequence data": S1 Text

S1 Text. *HvThf1* prey sequences.

*HvThf1* prey fragment

GACAGGGATGCCATCTTCAAGTCATATGTAACGGCGTTAAATGAAGATCCTGAGCAATACA
GGGCTGATGCACAAAGGATGGAAGAGTGGGCACGCTCACAAAATGGTAATCTGTTAGTTG
AATTTTCTTCCAGAGATGGGGAAATAGAGTCCATTCTGAAAGATATATCAGAAAGGGCCCA
GGGTAAGGGAAACTTCAGCTACAGCCGGTTCTTTGCTGTTGGCTTGTTCCGTTTGCTTGAG
CTCTCAAATGCGACGGAGCCAACCGTACTGGACAAGCTTTGCGCTGCTCTAAACATCAATA
AAAAAAGCGTGGATAGAGACCTTGACGTTTACCGCAACTTACTGTCGAAATTGGTTCAAGC
CAAGGAACTTCTTAAGGAATACGTCGAGAGGGAAAAGAAGAAGAGAGCAGAGAGATTGGA
GACGCCCAAGCCGAACGAGGCTGTTGCAAAATTCGATGGAAGCACTTATCCCTTGAAGCA
TTAACTCAACCTTTCCCCAGAGGAGAGCTTGTCCGGATAACAGATATTAACAGTTATACATC
TGGATAGCGTTGAGAACTATCGGGGGCATCTTTGGCTGCTCTTTGCGTGGTGTAGTACTCG
GCGGTACAAAAAGATCCAGATTATTCGCAGTAGTTGGTATTTTGTAATTTCTCCTTCTCTGT
GTTCGAATGTACCGTTTTCTTCACTTCGTTGACAATATATGAAGAGTTATCTGGTGGTGAAT
GTTACTCACAAGATGAGAAATCATATATTTTGGTGAAAAAGAAGTCATTCTGACTGGTGCTG
TTAAAAAAAAAAAAAAAAAAAA

HvTHF1 protein translation

DRDAIFKSYVTALNEDPEQYRADAQRMEEWARSQNGNLLVEFSSRDGEIESILKDISERAQGK
GNFSYSRFFAVGLFRLLELSNATEPTVLDKLCAALNINKKSVDRDLDVYRNLLSKLVQAKELLKE
YVEREKKKRAERLETPKPNEAVAKFDGSTYPLKH
